# Aurora B is required for programmed variations of cytokinesis during morphogenesis in the *C. elegans* embryo

**DOI:** 10.1101/319657

**Authors:** Xiaofei Bai, Po-Yi Lee, Chin-Yi Chen, James R. Simmons, Benjamin Nebenfuehr, Diana Mitchell, Lindsey R. Klebanow, Nicholas Mattson, Christopher G. Sorensen Turpin, Bi-Chang Chen, Eric Betzig, Joshua N. Bembenek

## Abstract

While cytokinesis has been intensely studied, how it is executed during development is not well understood, despite a long-standing appreciation that various aspects of cytokinesis vary across cell and tissue types. To address this, we investigated cytokinesis during the invariant *C. elegans* embryo lineage and found several reproducibly altered parameters at different stages. During early divisions, furrow ingression asymmetry and midbody inheritance is consistent, suggesting specific regulation of these events. During morphogenesis, we find several unexpected alterations including migration of midbodies to the apical surface during epithelial polarization in different tissues. Aurora B kinase, which is essential for several aspects of cytokinesis, remains localized to the apical membrane after internalization of other midbody components. Inactivation of Aurora B causes cytokinesis failure, which disrupts polarization and tissue formation. Therefore, cytokinesis shows surprising diversity during development and is required during epithelial polarization to establish cellular architecture during morphogenesis.

## Introduction

Generation of a multicellular organism requires that carefully orchestrated cell division is integrated properly into different developmental processes. Cell division is required not only to generate new cells that organize into tissues, but also to dictate the size, position and timing of daughter cells that are generated. Several aspects of cell division, including spindle orientation and division symmetry are well known instruments of developmental programs (Siller and Doe, 2009). Roles for cytokinesis in regulating developmental events are emerging, but are much less understood (Chen et al., 2013; Herszterg et al., 2014; Li, 2007). Using advanced live imaging, we sought to investigate cytokinesis in the well-defined divisions of the invariant *C. elegans* embryo lineage, which has been completely described (Sulston et al., 1983).

Cytokinesis is the final step of cell division and is normally a constitutive process during the exit from mitosis defined by discrete steps that occur during “C phase” (Canman et al., 2000; Oegema and Hyman, 2006). During cell division, signals from the anaphase spindle initiate ingression of the cleavage furrow (Bringmann and Hyman, 2005; Eggert et al., 2006), which constricts the plasma membrane onto the spindle midzone and leads to formation of the midbody. The midbody is a membrane channel between daughter cells containing microtubules and a defined organization of more than one hundred proteins that collaborate to execute abscission, the final separation of daughter cells (Green et al., 2012; Hu et al., 2012; Skop et al., 2004). Many of the proteins that contribute to midbody formation and function have roles in the formation of the central spindle and the contractile ring (El Amine et al., 2013). In addition, vesicles are delivered to the midbody that contribute lipids as well as regulators of abscission (Schiel et al., 2013). Subsequently, the ESCRT machinery assembles, microtubules are cleared and membrane scission occurs (Guizetti et al., 2011; Schiel et al., 2011). Aurora B kinase (AIR-2 in *C. elegans*) is essential for cytokinesis, acting on a number of targets although its precise mechanism is not fully elucidated. Aurora B also prevents completion of abscission in response to developmental or cell cycle cues partly by regulating the ESCRT machinery (Carlton et al., 2012; Carmena et al., 2015; Mathieu et al., 2013; Norden et al., 2006; Steigemann et al., 2009). How abscission is regulated in different developmental contexts has not been thoroughly investigated. In general, while mechanistic details of cytokinesis are being elucidated, it is generally assumed that these events occur through a standard, well-defined series of ordered events.

Exceptions to such a clear linear view of cytokinetic events have long been known, but are considered to be specialized cases. The most extreme examples are cells that do not complete cytokinesis altogether and become polyploid, such as liver or intestinal cells (Amini et al., 2015; Fox and Duronio, 2013; Hedgecock and White, 1985; Lacroix and Maddox, 2012). Another well-known example is found in several systems where germ cells do not complete abscission and remain connected through ring canals, which can allow flow of cytoplasm into germ cells (Greenbaum et al., 2007; Haglund et al., 2011; Hime et al., 1996; Maddox et al., 2005). Delayed abscission has also been observed in other cell types to keep daughter cells connected (McLean and Cooley, 2013; Zenker et al., 2017). Other variations of cytokinesis include cleavage furrow re-positioning during anaphase to change the size and fate of daughter cells (Ou et al., 2010). The symmetry of furrow ingression is important in established epithelial tissue where the furrow constricts toward the apical side of the cell and must occur while appropriate cellular contacts are preserved (Herszterg et al., 2014). In zebrafish neuroepithelial divisions, asymmetrical furrowing positions the midbody at the apical domain, which is inherited by the differentiating daughter (Paolini et al., 2015). Therefore, there are a number of ways the standard pattern of cytokinesis can be altered and more investigation is required to understand the functional purpose of these changes and how they are achieved.

Recent studies of abscission have sparked renewed interest in the midbody, which has led to insights into other functions it has beyond abscission (Chen et al., 2013). Recent evidence indicates that the midbody is cut off from each of the daughter cells that give rise to it (Crowell et al., 2014; Konig et al., 2017). The midbody may then be engulfed by either cell or persist extracellularly, which can depend on cell type (Ettinger et al., 2011; Salzmann et al., 2014). The midbody can also travel to non-parent cells, suggesting that it may carry or transport signals between cells (Crowell et al., 2014). The midbody is reproducibly inherited in *Drosophila* germline stem cells, but does not always end up in the stem cell (Salzmann et al., 2014). In dividing neuroepithelial cells, a stem cell marker is concentrated at the midbody and released into the lumen of the neural tube, which might provide signals during neuronal development (Dubreuil et al., 2007). This has led to the hypothesis that the midbody provides cues that regulate cell fate, although a detailed mechanistic understanding of this has not been elucidated.

A more clearly defined function for the midbody has been uncovered in cells that undergo polarization events after the completion of cytokinesis. For example, Madin-Darby canine kidney (MDCK) cells can establish apical basal polarity and organize into a simple epithelial lumen structure (Reinsch and Karsenti, 1994). Apical membrane markers are first delivered to the midbody during cytokinesis, establishing an apical membrane at the interface between the first two daughter cells (Schluter et al., 2009). Proper abscission and midbody positioning is required, in addition to proper spindle orientation, for MDCK lumen formation (Lujan et al., 2016; Reinsch and Karsenti, 1994). Polarized trafficking during cytokinesis has been shown to promote lumen formation in other systems as well (Wang et al., 2014b). Abscission is also delayed in acentrosomal blastomeres of the early mouse embryo to generate a midzone-derived microtubule organizing center that directs delivery of apical membrane markers to the plasma membrane (Zenker et al., 2017). The midbody becomes the apical process in chick neuronal progenitors (Wilcock et al., 2007) and defines the site of polarization for dendrite extension in *D. melanogaster* neurons (Pollarolo et al., 2011). The midbody is also a polarizing cue in the *C. elegans* embryo during the establishment of dorsoventral axis formation (Singh and Pohl, 2014; Waddle et al., 1994). In addition, the midbody can play a role in cilium formation (Bernabe-Rubio et al., 2016). Further effort is required to understand how cytokinesis and the midbody regulate pattern formation in tissues.

We sought to better understand how different aspects of cytokinesis are regulated during development by taking advantage of the well characterized invariant *C. elegans* embryonic lineage. We investigated furrow symmetry, central spindle dynamics, abscission timing and midbody inheritance. We find that cytokinesis follows a lineage specific pattern and that furrow symmetry and midbody inheritance is highly reproducible. During morphogenesis, abscission is delayed and the midbody forms in the middle of the cell and migrates to the apical membrane in the developing digestive and sensory tissues in *C. elegans*, which is a novel behavior. Unlike typical cytokinesis events, different midbody components have different fates in different tissues. Notably, Aurora B kinase remains localized to several apical surfaces after internalization of other ring components. Inactivation of temperature-sensitive Aurora B mutants causes sporadic cytokinesis defects that disrupt apical polarization of microtubules and adhesion. Cytokinesis mutants also have disrupted tissue organization, indicating an important role for specialized cytokinesis during morphogenesis. Together, our results suggest that Aurora B kinase has a novel function at the apical surface after cytokinesis during epithelial polarization and that modified implementation of cytokinesis is critical for proper cellular organization during animal development.

## Results

### Cytokinesis in the first two mitotic divisions: asymmetric midbody inheritance

We sought to systematically examine cytokinesis using lattice light sheet and spinning disc confocal microscopy during the stereotypical divisions of the *C. elegans* embryo, which has been extensively studied primarily in the first cell division due to its size and ease of access. The first mitotic division of the P0 cell generates the larger anterior daughter AB and the posterior daughter P1 (Fig. 1 A). We observed different components that allow us to evaluate specific aspects of the cytokinetic apparatus including the central spindle, the cytokinetic furrow and the midbody. We also chose midbody markers that localize to the flank and ring sub-structures of the midbody (Green et al., 2012). To observe the midbody flank region, we imaged Aurora B kinase (AIR-2), microtubules, and the membrane trafficking regulator RAB-11 (Fig. 1, 3, and Video S1). We also imaged midbody ring markers including the non-muscle myosin NMY-2, which also labels the contractile ring, and the centralspindlin component ZEN-4 (Fig. 1 G-P and Video S1). While the first mitotic furrow shows some variable asymmetry as previously reported (Maddox et al., 2007), the midbody forms in a relatively central position between daughter cells (Fig. 1 B-C, G-H and L-M). AIR-2::GFP, endogenous AIR-2 staining and tubulin show the expected pattern of localization on the central spindle and midbody as expected (Fig. 1 B-C, Fig. S1 A-D, Fig. 3 A-B and Video S1). The midbody from the first mitotic division is always inherited by the P1 daughter cell (Fig. 1 A). The midbody microtubule signal diminishes within 8 minutes after furrowing onset, which is a general indicator of abscission timing (Fig. 3 B, I) (Green et al., 2013; Konig et al., 2017). AIR-2 is lost from the flank over time but can be observed on the midbody remnant even after it is internalized into P1 (Fig. 1 D-E and Video S1). Additionally, each of the ring components behaves similarly to AIR-2, as expected (Fig. 1 I-J and N-O). Therefore, AIR-2 and other ring components remain co-localized on the midbody throughout the final stages of cytokinesis and are reproducibly internalized by the P1 daughter cell, as previously observed (Bembenek et al., 2013; Ou et al., 2014; Singh and Pohl, 2014).

**Figure 1.**
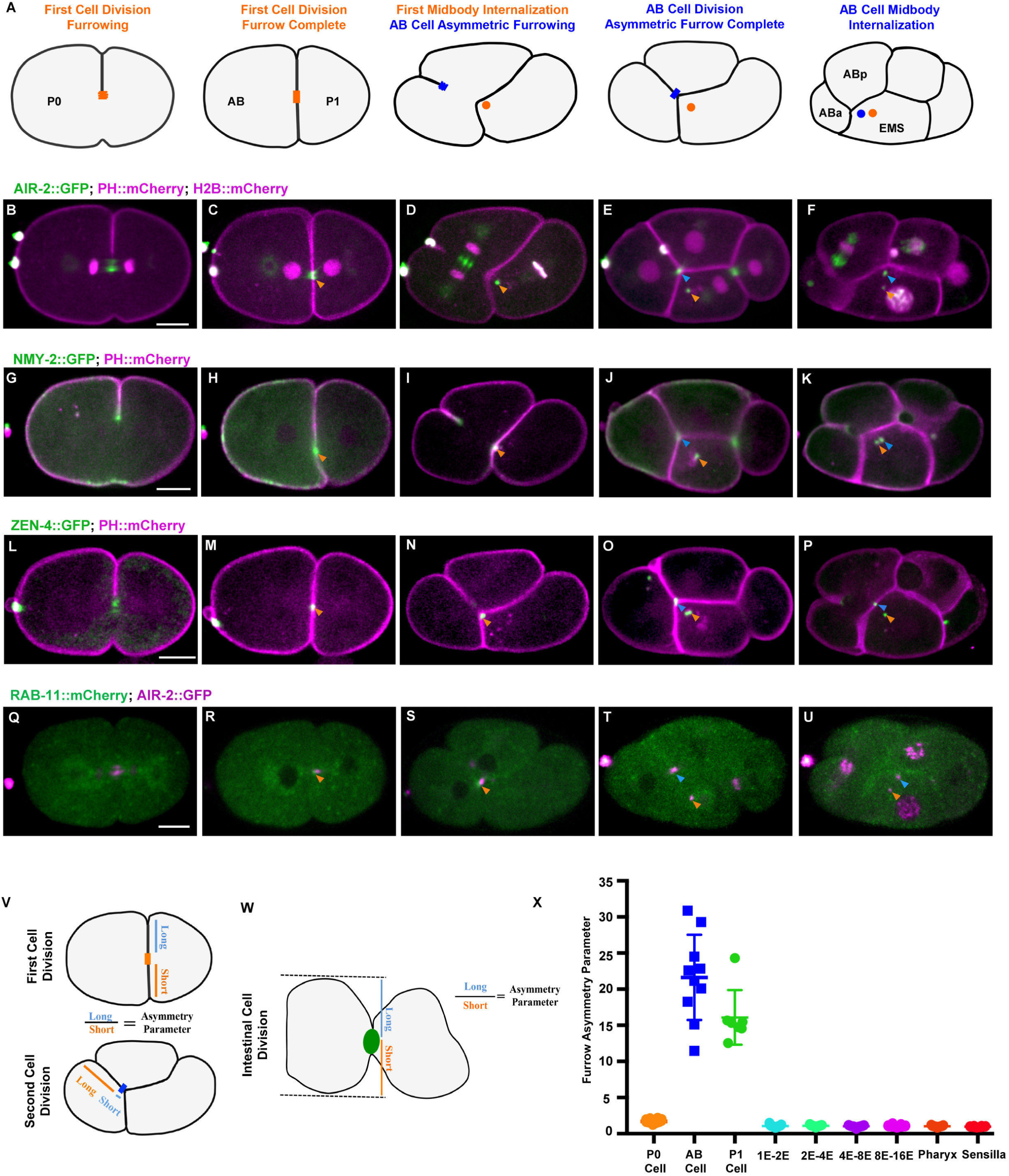
Cytokinesis in the first two mitotic divisions. (A) Illustration of cytokinesis in the first two mitotic divisions indicating the invariant fate of the midbody after division. Orange shapes indicate the first midbody, while blue shapes indicate the AB midbody. (B-F) Cytokinesis labeled with AIR-2::GFP (green, PH::mCherry and H2B::mCherry in magenta). During late anaphase, Aurora B localizes on the central spindle (B) which condenses into the midbody flank (C, orange arrowhead) and remains on the midbody until it is internalized by the AB daughter cell (D, orange arrowhead). During the second mitosis, the furrow is highly asymmetric and sweeps the central spindle against the EMS boundary, where the midbody forms (E, blue arrowhead). EMS engulfs the midbody instead of either of the AB daughter cells (F, blue arrowhead). (G-K) NMY-2::GFP (green, PH::mCherry in magenta) show localization to the furrow (G) and midbody ring (H-K). (L-P) ZEN-4::GFP (green, PH::mCherry in magenta) appears on the central spindle (L) and the midbody (M-P). (Q-U) RAB-11::mCherry (green) co-localized with AIR-2::GFP (magenta) briefly at the midbody, but does not remain on the midbody once it is internalized into cytosol (R-U). (V-X) Quantification of furrow asymmetry, measurement illustrated in V, W. (X) Furrow asymmetry parameter is significantly greater in second cell division. Scale bar, 10 μm.

During the second round of division, we observed substantial, reproducible changes in the pattern of cytokinesis, beginning with furrow symmetry. During the AB daughter cell division, which gives rise to ABa and ABp, the furrow ingressed from the outer surface until it reached the opposite plasma membrane in contact with EMS (Fig. 1 D-E, I-J, N-O). We calculated a symmetry parameter using the ratio of furrow ingression distance from each side of the furrow at completion (Maddox et al., 2007). On average, the furrow symmetry parameter is 1.7 in the first division, while the AB furrow is 21.6 and the P1 furrow is 16.1, indicating highly asymmetric furrows in the second divisions (Fig. 1 V, X). The central spindle is swept from the middle of the AB cell into contact with EMS during furrow ingression (Fig. 1 E, Video S1). AIR-2 localizes to the central spindle, then the midbody flank and remains associated with the midbody remnant after it is engulfed (Fig. 1 D-F, S-U and Video S1). NMY-2 and ZEN-4 also follow the expected pattern during cytokinesis and appear on the midbody that forms in contact with EMS (Fig. 1 I-J, N-O and Video S1). RAB-11::GFP accumulates briefly prior to abscission in both the first and second rounds of division and is not observed on the midbody afterward (Figure 1 Q-U), as shown previously (Ai et al., 2009; Bai and Bembenek, 2017; Bembenek et al., 2010). Therefore, all midbody markers examined behave as expected during the first two divisions.

Our examinations confirmed the unusual and consistent pattern of midbody inheritance after the AB division. The midbody from the AB cell division that forms in contact with EMS after highly asymmetric furrowing is invariably engulfed by EMS instead of either of the AB daughter cells (Fig. 1 F, K, P, U, Fig S1 D and Video S1). Further, the midbody from the P0 division is also always inherited by EMS. Microtubules in the midbody flank disappear within 8 minutes after furrowing in both AB and P1 cell divisions, indicative of relatively fast abscission at this stage (Fig. 3 C-D and I). This analysis confirms expected patterns of midbody regulation and midbody protein dynamics during the early embryonic divisions. These data reveal reproducible furrow ingression symmetry and midbody inheritance, which likely involve multiple mechanisms operating during cytokinesis in order to achieve this highly reproducible pattern.

### Aurora B localizes to the apical membrane after midbody migration and internalization in E8-E16 gut cytokinesis

We next performed a similar detailed analysis of cytokinesis on three developing tissues in morphogenesis, which revealed several novel patterns of cytokinesis. During morphogenesis, cells undergo terminal divisions and start to form tissues by polarizing and changing shape. The intestine is a well characterized epithelial tube derived from the E blastomere that undergoes five embryonic divisions (Leung et al., 1999). The E8 to E16 division occurs around 280 minutes after the first cleavage, after which cells undergo a mesenchyme to epithelial transition involving epithelial polarization and subsequently organize into a tube (Leung et al., 1999). Our observations demonstrate that these cells are performing a highly modified cytokinesis as they undergo polarization, which to our knowledge has not been previously reported (Fig. 2 A). The E8 cells undergo symmetrical furrowing to produce a centrally placed midbody (Fig. 2 B, E, I, Fig. S2 A and Video S2-4) with a 1.0 symmetry parameter (Fig 1 W, X), in contrast with the highly asymmetric furrow and displaced midbody location during the AB cell division. Therefore, the E8-E16 division appears largely routine up until the point of midbody formation.

**Figure 2.**
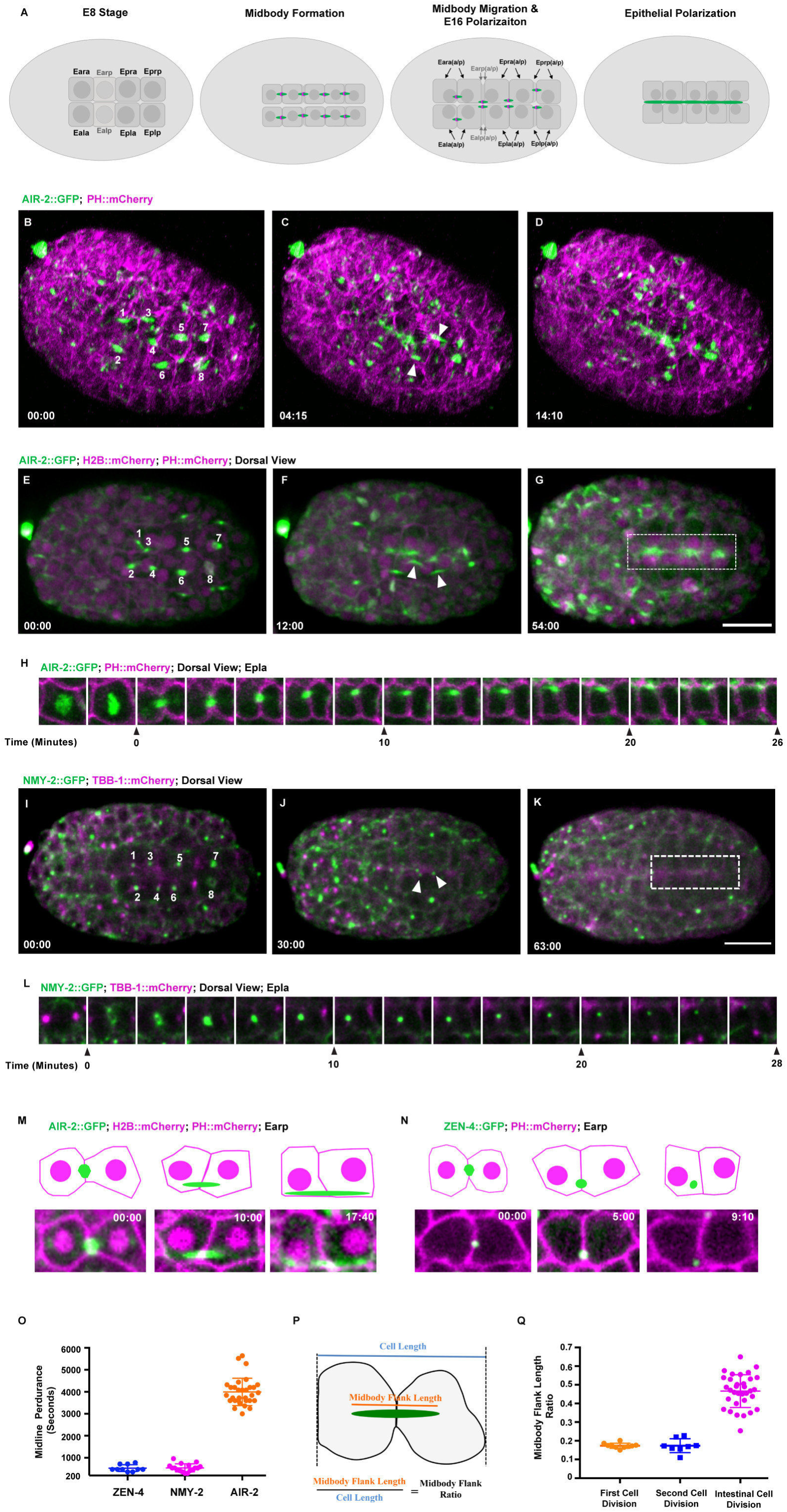
Midbody migration and Aurora B apical localization in the E8-E16 intestinal divisions. (A) Diagram of cytokinesis in the intestinal E8-E16 mitotic divisions indicating localization of Aurora B (green, midbody ring in magenta). (B-D) Lattice light sheet imaging of E8-E16 intestinal cell divisions in embryos expressing AIR-2::GFP (green) with PH::mCherry (magenta). AIR-2::GFP labels midbodies (labeled 1-8 in B) that form in the middle of E8 daughter cell pairs and then migrate (arrowheads, C) to the nascent apical membrane where it persists well after polarization is complete (D). (E-G) Spinning disc confocal microscopy of AIR-2::GFP (green) with H2B::mCherry and PH::mCherry (magenta) showing midbody migration (F) and apical localization (G). (H) Image series of Epla division with AIR-2::GFP (green, PH::mCherry in magenta) starting in prometaphase, clearly indicating midbody formation (t=0) and migration to apical midline. (I-K) NMY-2 (green, microtubules in magenta) localizes to furrows, then midbody rings (labeled 1-8 in F) that move to the midline (arrowheads, J) but do not persist (rectangle box in K). (L) Montage showing a single NMY-2::GFP labeled midbody migrating to midline. (M-N) Single plane imaging of midbody dynamics in individual intestine cell shows extension of the central spindle and apical membrane localization of AIR-2::GFP (M) and rapid internalization of ZEN-4::GFP (N) to the cytosol. (O) Quantification of midline perdurance of different midbody components (measured from the end of furrowing to internalization or loss of signal). (P) Illustration of E8 division and (Q) quantification of the ratio of maximal midbody flank length to cell length in different divisions. Scale bar, 10 μm. Time shown in minutes:seconds. Error bars indicate standard deviation of the mean.

After midbody formation, there are several changes to the pattern of cytokinesis, which occur during epithelial polarization. Using lattice light sheet imaging, we observe that centrally located midbodies from both left and right daughter cell divisions (Ealp, Earp, Epla and Epra) migrate across the width of the cell to the nascent apical surface at the midline, which completes 30 minutes after furrow ingression (Fig. 2 B-D and Video S2). The midbody flank region elongates during the migration process and the flank microtubules persist for over 25 minutes on average from furrow ingression to when they join other microtubules at the apical midline and can no longer be distinguished, which is three times longer than the early divisions (Fig. 3 F, H-I). AIR-2::GFP localizes along the extended length of the spindle midzone microtubules as they move to the apical midline instead of becoming confined to the midbody remnant as observed in early divisions (Fig. 2 E-H, M, and Video S2-4). The ratio of the length of this midbody flank relative to the cell at the greatest length is 0.47 (average 4.6 μm / 9.8 μm) in the intestinal cell division, which is more than twice that of the early two cell divisions 0.17 (average 9.3 μm/ 53.4 μm) in P0 and 0.17 (average 7.7 μm/ 44.3 μm) in AB (Fig. 2 P-Q). The ring markers ZEN-4 and NMY-2 are quickly internalized (553±140 seconds and 545±179 seconds, respectively) after the midbody reaches the apical midline (Fig. 2 I-L, N-O and Video S3). Collectively, these data indicate that abscission occurs after migration of the midbody to the apical midline. Therefore, E8 cells undergo a programmed apical midbody migration event instead of having an asymmetrical furrow lead to the formation of an apically localized midbody, as observed in the AB cell division and epithelial cells in other systems. To our knowledge this is the first observation of collective midbody migration and may represent a new stage of cytokinesis.

In addition to this midbody migration event, we observed a novel behavior of AIR-2 during E8-E16 cytokinesis. In contrast to the midbody ring components, AIR-2 remains localized at the apical midline well after the time that ring components are internalized and polarization is complete (Fig. 2 D, G, H, M and Video S2-4), co-localizing with the apical polarity marker PAR-6 (Fig S2 F-H). Endogenous AIR-2 can also be observed at the apical midline by immunofluorescence (Fig. S1 E-G). The gut apical surface recruits pericentriolar material donated by the centrosome during E16 polarization (Feldman and Priess, 2012; Yang and Feldman, 2015). We observed that γ-tubulin::GFP moves to the apical surface at the same time as AIR-2::GFP (Fig. S2 I). High temporal resolution single plane confocal imaging tracking individual midbody dynamics confirm the elongated AIR-2::GFP flank localization and persistence at the apical midline compared to the rapid internalization of ZEN-4::GFP after the migration event (Fig. 2 M-N and Video S4). For comparison, we imaged the first three E cell divisions and found symmetric furrowing (Fig. 1X), rapid abscission timing (Fig. 3I) and that midbody behavior was similar to the early embryo divisions with no exaggerated alterations (Fig. S3). Therefore, different midbody components have different fates at the end of cytokinesis specifically in the E8-E16 intestinal divisions during epithelial polarization, with ring markers being internalized while AIR-2 remains at the apical surface. To our knowledge, this is the first report of AIR-2 localization remaining at the plasma membrane after abscission.

**Figure 3.**
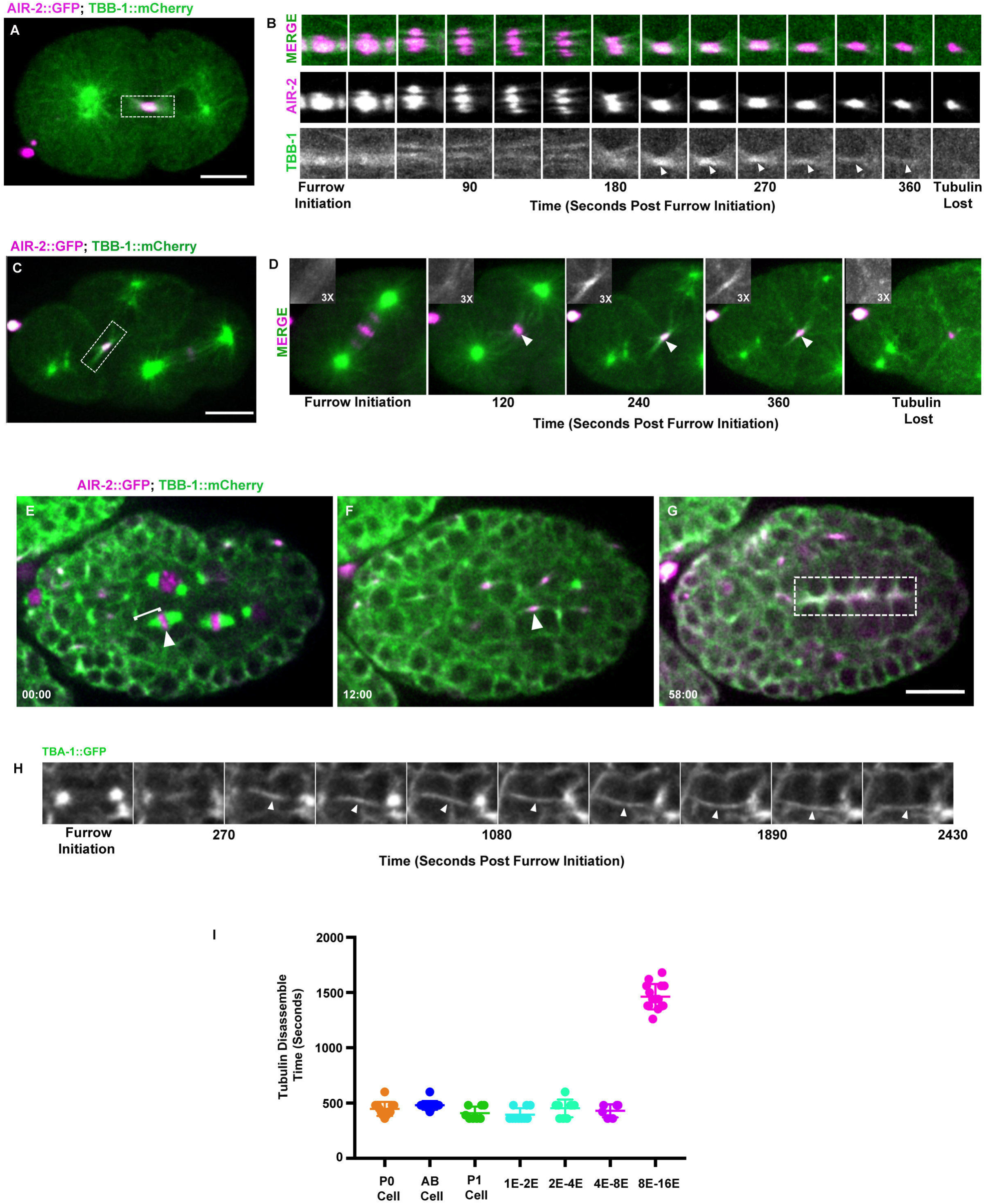
Abscission is delayed during gut epithelial polarization. Spindle midzone microtubule dynamics were imaged during different cell divisions to assess abscission timing. (A-B) AIR-2::GFP (magenta) and β-tubulin TBB-1::mCherry (green) colocalize at the central spindle during anaphase and furrowing in the first cell division. AIR-2::GFP persists at the midbody after microtubules are lost, which indicates rapid abscission timing (B). (C-D) AIR-2::GFP (magenta) and β-tubulin TBB-1::mCherry (green) colocalize on the central spindle of the AB cell that is swept adjacent to EMS after highly asymmetric furrowing. Microtubules are lost quickly like the first division (D). (E-G) Tubulin TBB-1::mCherry (green) and AIR-2::GFP (magenta) localize to an extended flank region during intestinal midbody migration (F, arrowhead). (G) Tubulin and AIR-2 persist at the apical membrane after polarization (rectangle box). (H) Single z-plane imaging of midbody flank microtubules (arrowheads) during Epra cell division and midbody migration. The extended flanking microtubules persist at least 3 times longer than earlier divisions throughout the migration process until they can no longer be distinguished from other microtubules at the apical surface. (I) Quantification of tubulin persistence time at the central spindle during different cell divisions. Scale bar, 10 μm. Error bars indicated standard deviation of the mean.

In other lumen forming systems, such as MDCK cells, RAB-11 vesicle trafficking during cytokinesis transports apical membrane components to the midbody to establish the apical membrane (Schluter et al., 2009). In *C. elegans*, RAB-11 endosomes control trafficking at the apical surface of the intestine throughout the life of the animal (Sato et al., 2014). We imaged RAB-11 during the E8-E16 division to examine when apical localization occurs. RAB-11::mCherry colocalizes with AIR-2::GFP once the midbody is formed and migrates to the apical surface with the midbody (Fig. S2 J-L). RAB-11::mCherry is also localized at spindle poles, as in other mitotic cells (Albertson et al., 2005), which also migrate to the apical surface (Feldman and Priess, 2012). Similar to AIR-2, RAB-11 remains localized to the apical surface well after cytokinesis is complete and remains at this position throughout the life of the animal (Fig. S2 L). These observations indicate that the apical localization of RAB-11 is established during cytokinesis in the E8-E16 division, which may contribute to formation of the apical surface during intestinal epithelial polarization.

The anterior and posterior pair of E16 cells (Ealaa, Earaa, Eplpp and Eprpp) undergo one last embryonic division to achieve the E20 intestine stage. In the four central E8 cells that do not divide again, the midbody migrates to the midline at E8-E16 as described above. However, the midbodies from the other four E8 cells (Eala, Eara, Eplp and Eprp), which undergo another division, migrate toward the midline but the AIR-2 signal diminishes (Fig. S2 E). During the final E16-E20 divisions, the midbodies of Ealaa, Earaa, Eplpp and Eprpp undergo apical migration after symmetrical furrowing (Fig. S2 M-O and Video S5). Therefore, the midbody migration event in the intestine does not happen only during the polarization event that occurs during E8-E16, suggesting that it is specifically programmed to occur during the terminal embryonic divisions. Post-embryonic divisions in the intestine at L1 lethargus involve nuclear but not cytoplasmic divisions leading to the formation of binucleate cells that subsequently undergo multiple rounds of endoreduplication to become highly polyploid (Hedgecock and White, 1985). Therefore, cytokinesis in the intestinal lineage undergoes distinct regulatory phases at different stages of development.

### Aurora B Kinase is required for normal E8-E16 cytokinesis and proper epithelial polarization

Given the pattern of cytokinesis during the E8-E16 division and the localization of AIR-2::GFP to apical structures, we sought to investigate Aurora B function at this stage of development. We first asked if cytokinesis is essential for the final stages of embryonic development since it is possible that morphogenesis could occur despite the presence of binucleate cells. To bypass the essential function of cytokinetic regulators during the early embryonic cell divisions, we inactivated temperature sensitive (ts) mutants after isolation of two-cell embryos at the permissive temperature (15 °C) and shifted them to non-permissive temperature (26 °C) at different embryo stages until they hatched. The *air-2(or207)* embryos have only 53.6% (37/69) hatching even when left at 15 °C through hatching, indicating that this mutant is sick even at permissive temperature, while wild-type N2, *zen-4(or153)* and *spd-1(oj5)* embryos are 100% viable when kept at 15 °C (Table 1). Embryos shifted to 26 °C after 4.5 hours (corresponding to late E4 to early E8 stages) or 6.5 hours (corresponding to E8-E16 stages) at 15 °C showed significantly increased lethality in both *air-2(or207)* and *zen-4(or153)*, but not *spd-1(oj5)* (Table 1), which correlates with the amount of cytokinesis failure observed. The few animals that were able to hatch in *air-2(or207)* and *zen-4(or153)* mutants had severe morphogenesis defects (data not shown). Mutant embryos shifted after the completion of all the developmental divisions at the comma to 1.5-fold stage were largely rescued for lethality and hatched at a rate similar to permissive temperature (Table 1). Therefore, these results are consistent with the hypothesis that cytokinesis is essential for the final stages of embryonic development during morphogenesis.

**Table 1.** Hatch rate of temperature sensitive mutants dissected at the two cell stage.

| Stage Before Shifting | Genotype | Hatch Rate % (Hatch Embryos/Total) |
| --- | --- | --- |
| 15 °C Forever | N2 | 100% (32/32) |
|  | *air-2(or207)* | 53.6% (37/69) |
|  | *zen-4(or153)* | 100% (28/28) |
|  | *spd-1(oj5)* | 100% (35/35) |
| E4-E8 | N2 | 100% (26/26) |
|  | *air-2(or207)* | 6.3% (2/32) |
|  | *zen-4(or153)* | 0% (0/57) |
|  | *spd-1(oj5)* | 100% (48/48) |
| E8-E16 | N2 | 100% (45/45) |
|  | *air-2(or207)* | 14.4% (13/90) |
|  | *zen-4(or153)* | 10.1% (10/99) |
|  | *spd-1(oj5)* | 100% (83/83) |
| Comma-1.5 Fold | N2 | 100% (36/36) |
|  | *air-2(or207)* | 33.7% (31/92) |
|  | *zen-4(or153)* | 85.7% (54/63) |
|  | *spd-1(oj5)* | 100% (27/27) |

Next, we tested whether Aurora B kinase was required for the specialized E8-E16 divisions and epithelial polarization. The *air-2(or207)* mutant has a high percent cytokinesis failure within minutes after shifting to non-permissive temperature in one-cell embryos (Severson et al., 2000), but does not show penetrant cytokinesis failures unless shifted several hours in older embryos (Figure 4A). We first asked whether microtubule organization at the midbody and apical surface required AIR-2 function. Live cell imaging of *air-2(or207)* mutants expressing TBB-1::GFP shifted at the E4-E8 stage demonstrated that spindle midzone microtubules were reduced relative to wild-type E8 cells (Fig. 4 B-I, Video S6). Inactivation of *air-2(or207)* under these conditions caused 27% of the observed E8 cells to fail cytokinesis, which causes these cells to fail to polarize all nuclei at the midline (Fig. 4F-L). In cells that do not fail cytokinesis, weak spindle midzone microtubules can be observed, which move to the apical surface where microtubules quickly accumulate similar to WT (Fig. 4 B-I, Video S6). In neighboring cells that fail cytokinesis, microtubule accumulation at the apical midline is disrupted, which is most obvious when both left and right E8 divisions fail at the same time (Fig. 4 H, J, K, Video S6). Therefore, Aurora B kinase is required for central spindle dynamics and completion of cytokinesis during the E8-E16 divisions, which is required for normal nuclear polarization and accumulation of microtubules at the apical midline.

**Figure 4.**
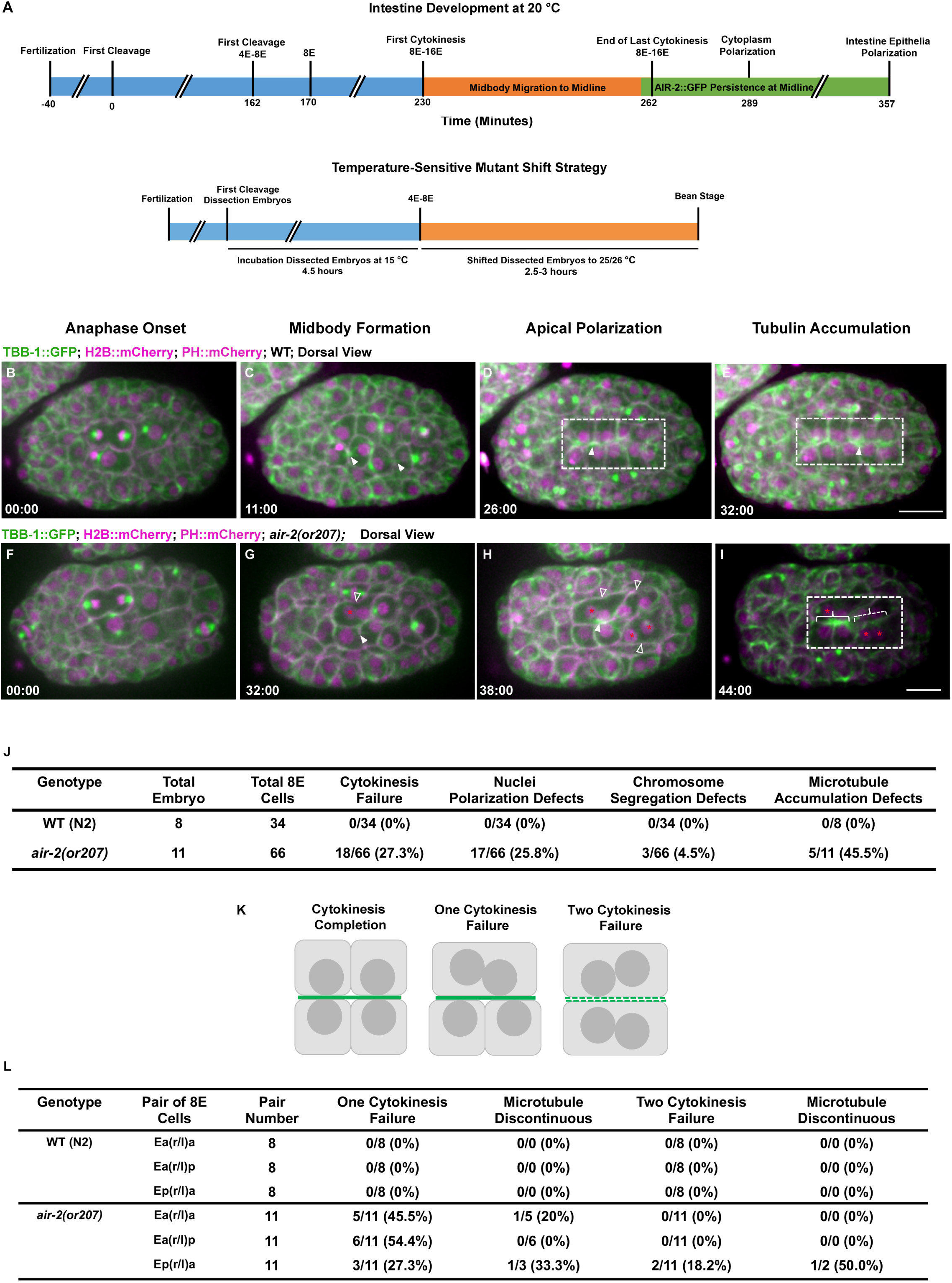
Aurora B is required for E8-E16 cytokinesis and epithelial polarization. Timeline of cell division events in the intestine (upper timeline) and the temperature shift strategy inactivating mutants from the end of the E4-E8 division through E8-E16 cytokinesis (A). (B-E) Imaging of microtubules (TBB-1::GFP, green) together with H2B::mCherry and PH::mCherry (magenta) shows spindle midzone microtubules (arrows, C) that migrate to the apical midline (D) where microtubules accumulate after polarization (E). In Aurora B mutant embryos (F-I), spindle midzone microtubules are diminished (G, solid arrowhead). Frequent cytokinesis failures occur (G, H, open arrowheads) and nuclei fail to migrate to the apical midline (G-I, red asterisks). (J) Quantification of defects in Aurora B mutant E8-E16 divisions showing significant cytokinesis failures where most cells that failed also have nuclear polarization defects. Of the 11 embryos observed, five have apical microtubule accumulation defects. The low incidence of chromosome segregation defects indicate weak Aurora B inactivation compared with early embryo divisions in this mutant. (K) Diagram of failure patterns of pairs of E8-E16 divisions on opposite sides of midline and observed consequence on microtubule accumulation (green) at apical midline. (L) Quantification of individual cell division failures and incidence of discontinuous microtubule accumulation at the apical midline after polarization. Scale bar, 10 μm.

To more clearly observe formation of the apical surface, we imaged the dynamics of the adhesion complex during the E8-E16 divisions, which accumulates at the apical surface during polarization to promote gut lumen formation (Achilleos et al., 2010). We imaged the α-catenin homologue, HMP-1::GFP, a cytoplasmic component of the cadherin-catenin adhesion complex that links cell-cell contacts with the actin cytoskeleton. Previously, it was shown that polarity proteins recruit adhesion complexes to cortical foci in the polar cortex (Achilleos et al., 2010), which associates with centrosomes and moves with them to the apical midline (Feldman and Priess, 2012). We observed that HMP-1::GFP is also localized to the furrow from the onset of furrow ingression, accumulates adjacent to spindle midzone microtubules and moves together with the midbody to the apical surface (Fig. 5A-E, Movie S7). Analysis of the first three E cell divisions also revealed localization of HMP-1::GFP to the furrow and membrane adjacent to the midbody, suggesting a dynamic interplay between adhesion and cytokinesis during the embryonic divisions (Fig. S4). In *air-2(or207)* mutants, furrow HMP-1::GFP was reduced during cytokinesis (Fig. 5 F-J, Video S7). In E8 *air-2(or207)* divisions that complete cytokinesis, HMP-1::GFP signal normally accumulated at the apical midline (Fig. 5 F-J, Video S7). However, in *air-2(or207)* E8 divisions that failed cytokinesis, accumulation of HMP-1::GFP was significantly delayed especially when pairs of E8 daughters on opposite sides of the midline both failed (Fig. 5 F-J, Video S7). Therefore, inactivation of Aurora B kinase disrupts normal α-catenin dynamics during cytokinesis and delays accumulation of α-catenin at the apical surface during polarization.

**Figure 5.**
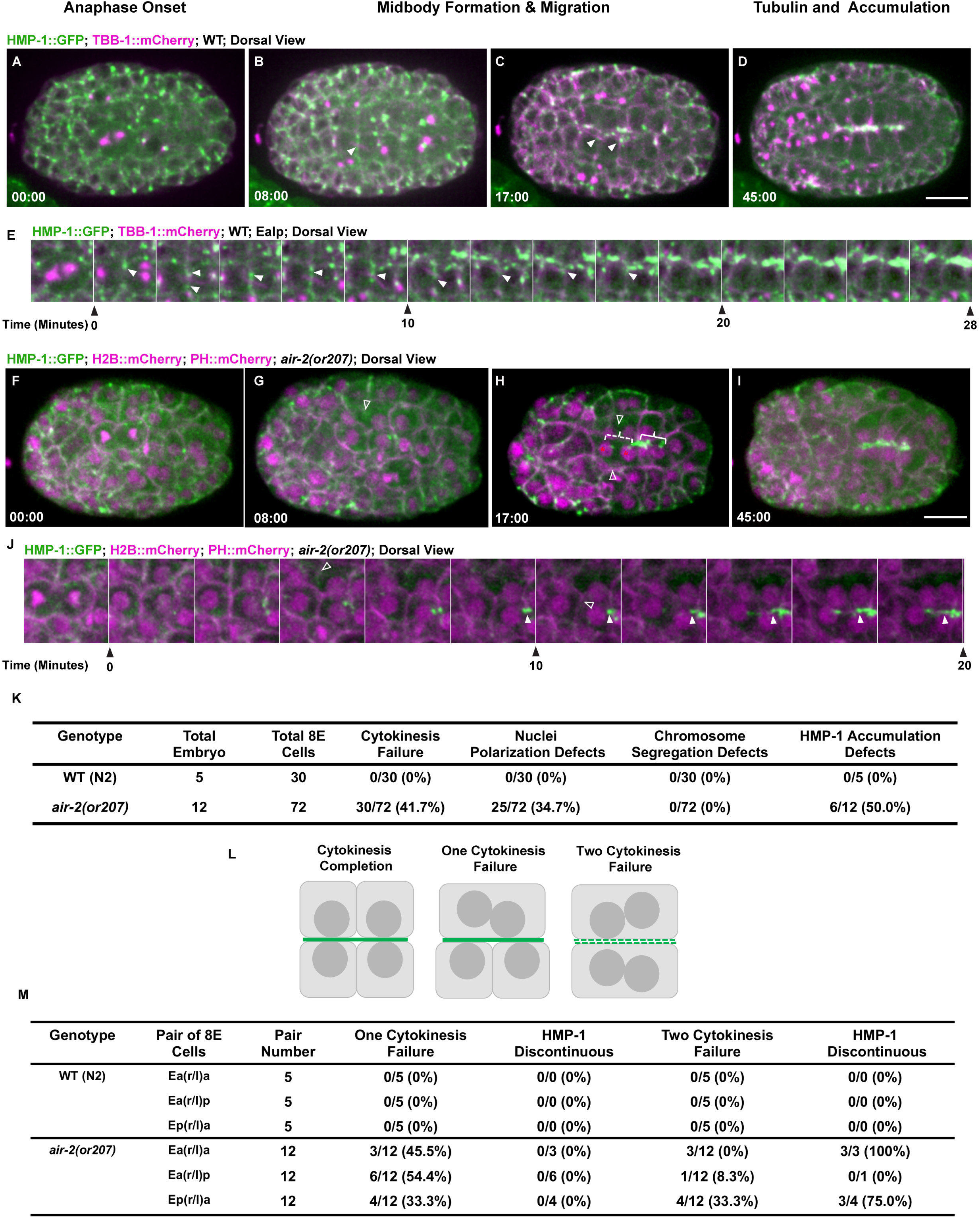
Aurora B is required for adhesion dynamics and timely apical adhesion accumulation during E8-E16 gut cytokinesis. Adhesion dynamics during E8-E16 intestinal division and polarization. (A-D) HMP-1::GFP (α-catenin, green, microtubules in magenta) localizes to the furrow and membrane adjacent to the midzone microtubules in the midbody (B, arrowhead) during cytokinesis. HMP-1::GFP localizes with the midbody as it migrates to the apical surface where it accumulates (C, D). (E) High temporal resolution imaging of HMP-1::GFP shows furrow and midbody accumulation (arrowheads) during cytokinesis and localization adjacent to midzone microtubules as they migrate to the apical midline. (F-I) Aurora B mutants disrupt adhesion dynamics. HMP-1::GFP is reduced on the furrow and midbody (F,G, open arrowhead). When both cells fail cytokinesis (G, H, open arrowheads indicate failures) on opposite sides of the midline, HMP-1 accumulation is delayed (dashed bracket indicates midline between two failed E8 gut cells, solid bracket indicates successful E8 division with apical accumulation). (I) HMP-1 signal eventually accumulates along the midline. (J) Kymograph of HMP-1::GFP in Aurora B mutant E8 cells that fail cytokinesis (open arrowheads indicate furrow regression) shows reduced localization to the furrow, midbody and delayed apical accumulation. (K) Quantification of defects in Aurora B mutant E8-E16 divisions showing significant defects in HMP-1::GFP accumulation with cytokinesis failures. (L) Diagram of cytokinesis failures in pairs of E8-E16 divisions on opposite sides of midline and observed consequence on adhesion accumulation (green) at apical midline. (M) Quantification of individual cell division failures and incidence of discontinuous adhesion accumulation at the apical midline after polarization. Scale bar, 10 μm.

In order to understand the effect of cytokinesis failure on lumen formation, we performed staining of the polarized gut using several apical markers. We shifted mutant embryos around the transition between the E4-E8 stages to 26 °C and fixed at the bean stage after intestinal polarization. We evaluated the formation of the apical surface of the intestine by staining for the Ezrin-Radixin-Moesin homologue, ERM-1 (van Furden et al., 2004). We stained *air-2(or207), zen-4(or153)* and *spd-1(oj5)* under these conditions and observed penetrant cytokinesis defects in *air-2(or207*) and *zen-4(or153)* but not *spd-1(oj5)* mutants (Fig. 6). In all cases, ERM-1 was localized to the apical surface of the intestine and pharynx (Fig. 6 A-F). However, ERM-1 staining was broadened, branched and/or discontinuous in *air-2(or207)* embryos (Fig. 6 B-D, H). Disrupted ERM-1 staining was also observed in *air-2(or207)* embryos, which were shifted at E4-E8 for 4.5-5 hours until the comma stage, indicating that these defects are not resolved later in development (Fig. S5 B, D). Furthermore, the intestine was highly mispositioned within the embryo as revealed by color-coded max Z-projections (Figure 6 H) and the nuclei were often randomly positioned on the z-axis compared with wild-type (Fig. 6 D-E inserts). The localization of other apical markers, such as PAR-3, DLG-1 and IFB-2, were similarly disrupted, localizing to the apical surface but in a disorganized way (Fig. S5 E-J). The *zen-4(or153)* embryos also had highly penetrant branched and discontinuous apical ERM-1 staining that was mispositioned within the embryo at a lower rate than the *air-2(or207)* mutants (Fig. 6 E, H and Fig. S5 C-D). The *spd-1(oj5)* embryos displayed a significant, but lower rate and severity of lumen defects (Fig. 6 F-H) despite having no lethality (Table 1) and minimal cytokinesis failures. Therefore, we conclude that proper execution of cytokinesis is required for normal lumen formation in the gut.

**Figure 6.**
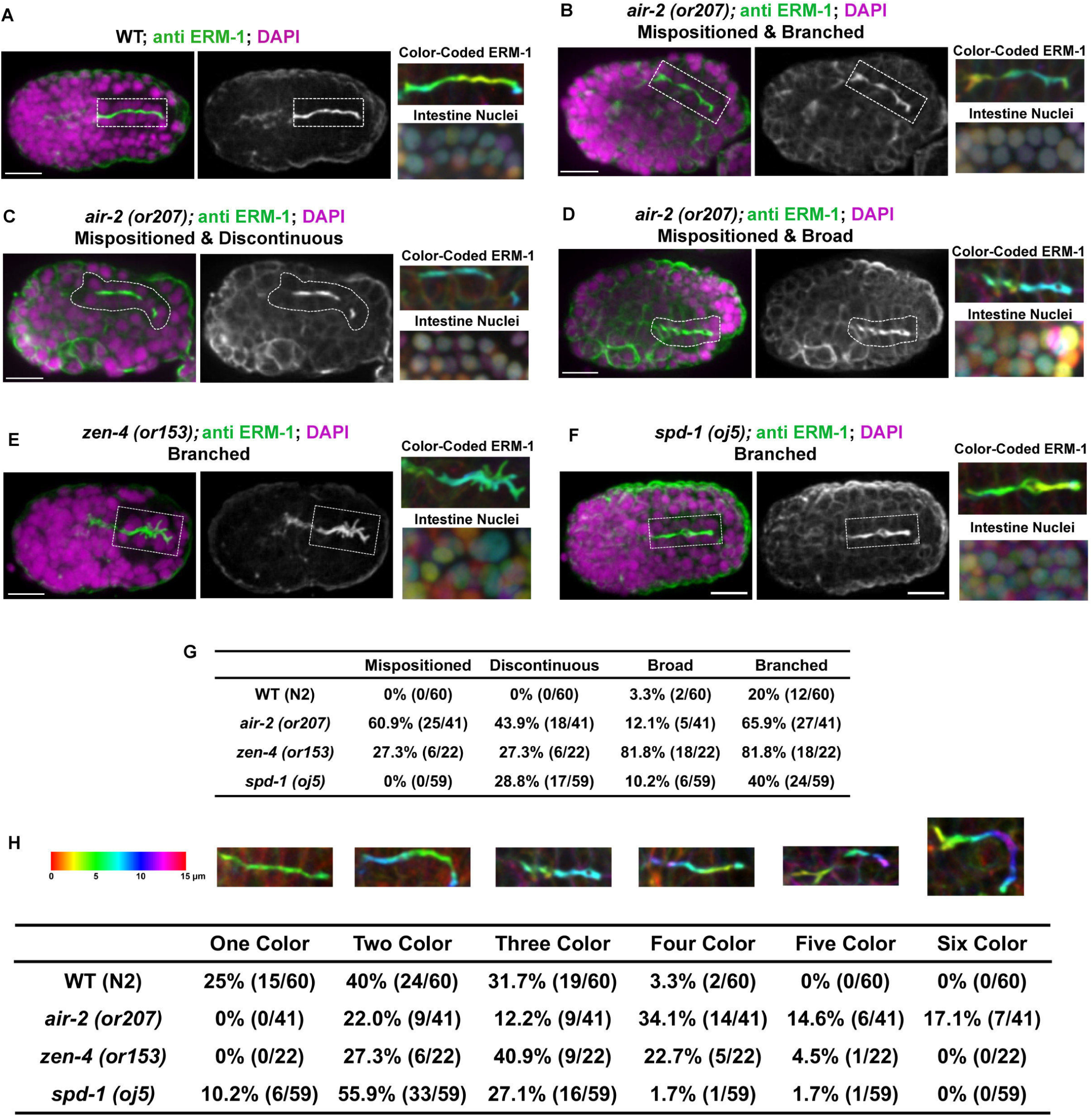
Gut morphogenesis is disrupted in cytokinesis mutants. Staining of apical markers reveals gut morphology after E8-E16 division and polarization. (A) ERM-1 staining in wild-type bean stage embryos is enriched at the apical midline of the intestine (dotted rectangle). Maximum z-projected images color-coded according to Z-depth (using FIJI temporal-color code plugin, scale shown in F) to visualize the three-dimensional position of ERM-1 and nuclei, which are mostly in a narrow range of Z-depth (A-F, H). In *air-2(or207)*, multiple defects are observed, including mispositioning of the entire intestine (B-D), branches in the apical surface (B), gaps in the apical surface creating a discontinuous lumen (C), or broader staining of ERM-1 (D). (E) The *zen-4(or153)* mutant causes many of these phenotypes, including branching of the apical surface and randomly positioned nuclei. (F) Subtle lumen and nuclei position defects are observed in *spd-1(oj5)* embryos. (G) Quantification of apical defects observed by ERM-1 staining in the different mutants. (H) Quantification of the defective z-plane distribution of the apical surface in the different mutants. The more colors a lumen has in the projection, the more skewed in the Z-axis it is within the embryo. Scale bar, 10 μm.

### Apical midbody migration and AIR-2 apical localization in the pharynx

We also noticed migration of the midbody and accumulation of midbody proteins at the apical surface during the terminal divisions in the pharynx. Unlike the intestine, which originates from a single blastomere that undergoes a very well defined series of divisions, the pharynx has a more complicated structure, containing more than 80 pharyngeal precursor cells (PPCs) that arise from both AB and MS founder cells (Sulston et al., 1983). The PPCs organize into a double plate structure prior to the final division, which occurs at around 310-325 minutes after the first cleavage, and then polarize and undergo apical constriction to become wedge shaped cells that form a lumen by 355 minutes (Rasmussen et al., 2013; Rasmussen et al., 2012). To obtain optimal images of this large, complex structure, we filmed at least a 15-micron Z-depth section of the embryo from both dorsal and ventral aspects with confocal microscopy (Figure 7, Video S8). We also filmed whole embryos with lattice light sheet microscopy, which provides higher spatial resolution during the pharyngeal cell division (Video S9). Similar to our observations in the intestine, PPCs are in the final stages of cell division as they polarize, which has not been previously described. PPCs undergo a symmetric furrowing event that yields a centrally placed midbody between the two daughter cells (Fig. 7A and Video S8). Also similar to the intestine, PPC midbodies migrate from their central position between daughter cells to the apical midline of the forming pharyngeal bulb (Fig. 7 F, K, O and Video S9). In PPC terminal divisions, AIR-2::GFP appears as a midbody flank structure that migrates to the apical midline and persists at the apical surface after cyst formation (Fig. 7 B-F and Video S9). ZEN-4::GFP appears on midbodies, migrates to the apical surface, and is rapidly degraded, similar to the intestinal divisions (Fig. S6 A-E and Video S8). NMY-2::GFP also labels midbodies and moves to the apical surface, but remains localized to the apical surface during apical constriction (Fig. 7 G-K). Similar to AIR-2, RAB-11 and tubulin accumulate and remain localized to the apical surface after polarization (Fig. S6 F-K). We confirmed endogenous AIR-2 localization with staining and show that it can be observed on the apical surface of the pharynx (Fig. S1 H-J). AIR-2 partially co-localized with PAR-6 at the apical membrane, which also acquires γ-tubulin::GFP after polarization (Fig. S6 L-Q). Also similar to the intestinal divisions, α-catenin (HMP-1::GFP) localizes to the furrow and adjacent to the midbody as it migrates to the apical midline where it accumulates after polarization (Fig. 7 L-O). Cytokinesis in the gut and pharynx show similar patterns where midbodies migrate to the apical midline and specific midbody components, especially AIR-2, remain localized at the apical cortex even after the midbody ring is removed. Therefore, similar patterns of apical localization and midbody migration are observed during epithelial polarization in the intestine and pharynx in *C. elegans*.

**Figure 7.**
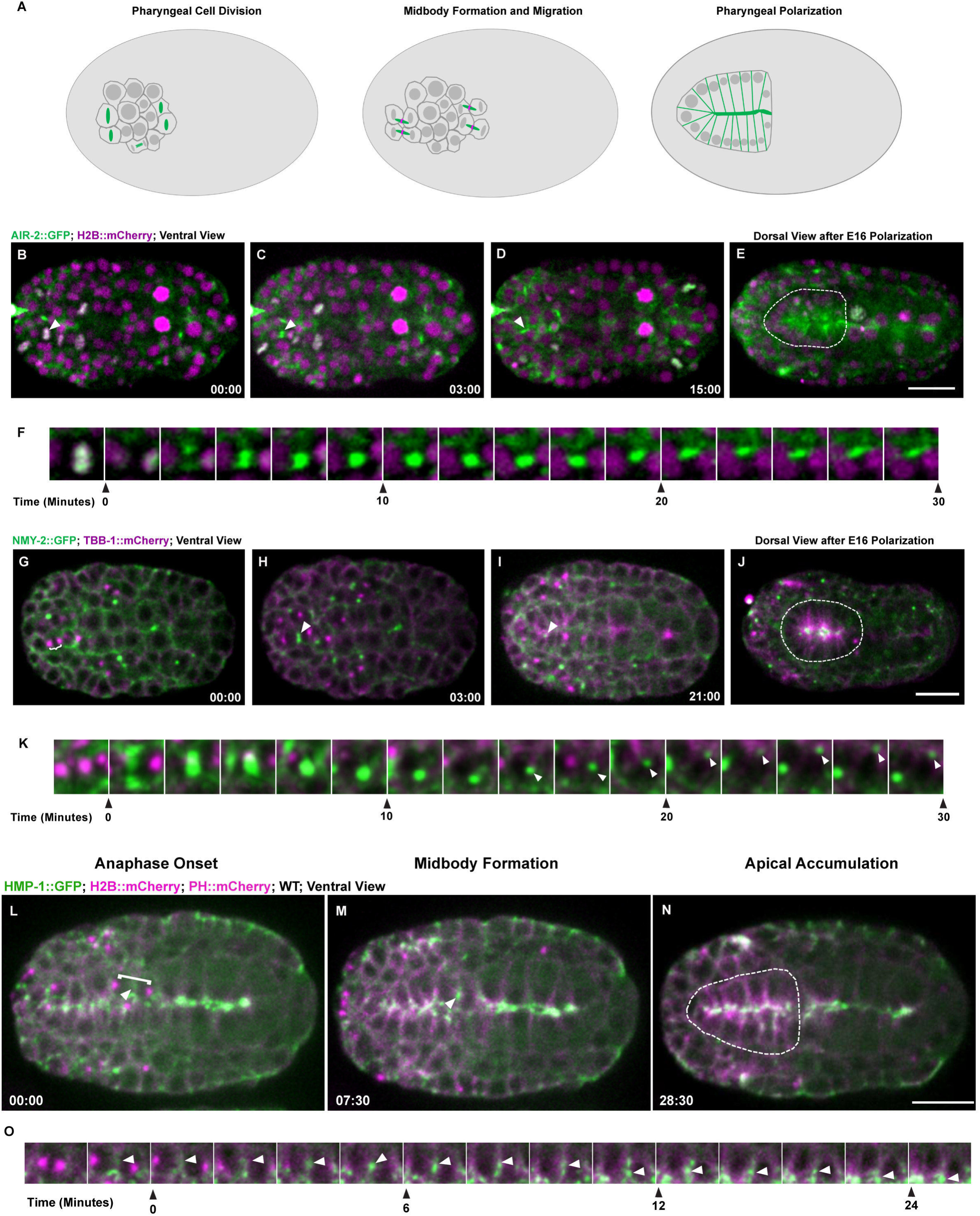
Cytokinesis During Pharyngeal Precursor Cell Polarization. (A) Illustration of the mesenchymal to epithelial transition of pharyngeal precursor cells (PPCs) showing cell division and dynamics of Aurora B (green, midbody ring in magenta). (B-E) PPC division labeled with AIR-2::GFP (green, H2B::mCherry in magenta) from both ventral (B-D) and dorsal (E) views. AIR-2::GFP localizes to chromatin in metaphase (B) and moves to the central spindle in anaphase (C) and appears on the midbody which moves toward the midline (D). AIR-2 persists at the apical surface for an extended time (E). (F) Image series showing an AIR-2::GFP labeled midbody migrating toward the midline. (G-J) Imaging NMY-2 (green, TBB-1::mCherry in magenta), shows the movement of the midbody to the midline (I, K). NMY-2 accumulates at the apical midline during apical constriction (J). (L-N) During cytokinesis, α-catenin (HMP-1::GFP, green, tubulin in magenta) accumulates on the furrow (arrowhead in L) and adjacent to the midbody (arrowhead, M) before accumulating at the midline (N). (O) Shows kymograph of HMP-1::GFP at the furrow and midbody, which migrates to the apical midline where HMP-1::GFP accumulates. Time shown in minutes: seconds. Scale bar, 10 μm.

### Apical midbody clustering and Aurora B dendrite localization in sensilla neurons

Finally, we observed a unique form of cytokinesis during the final divisions of the cells that make up the sensory neurons at the tip of the embryo. The *C. elegans* amphid sensilla are a sensory organ that contains 12 neurons with dendrites that extend processes through two sheath cells and the cuticle at the tip of the mouth. During morphogenesis, amphid neurons bundle together, anchor at the tip of the animal and migrate back to extend dendrites (Heiman and Shaham, 2009). From the lineage of the 12 sensilla neurons, there are 10 precursor cell divisions that occur between 280 and 400 minutes after the first cleavage (Sulston et al., 1983). These terminal divisions include two daughter cell pairs (ADF/AWB and ASG/AWA) and several where one daughter differentiates into a sensilla neuron while the other daughter undergoes apoptosis (ADL, ASE, ASK, ASI), or differentiates into another neuron (AWC, ASH, AFD, ASJ). Our observations show that these cells undergo a unique form of cytokinesis just before they undergo dendrite morphogenesis (Fig. 8 A). These cells undergo a symmetrical furrowing event before midbodies form centrally between the daughter cells (Fig. 8 B and Video S10-11). A group of at least 6 daughter cell pairs divide initially forming multiple midbodies as observed with both confocal and lattice light sheet imaging (Fig. 8 C and Video S10-11). These midbodies migrate together into a cluster over a 60-minute time window (Fig. 8 D). AIR-2::GFP, RAB-11 and tubulin persist in these clusters (Fig. 8 D, Fig. S7 A-F), while ZEN-4::GFP rapidly disappears during the midbody clustering process (Fig. 8 E and Video S10). Endogenous AIR-2 can detected by staining in these lateral apical clusters (Fig. S1 K-M). In contrast to ZEN-4::GFP, NMY-2::GFP migrates with the midbody to the cluster and persists at the very tip of the dendrites (Fig. 8 F and Video S10). We observe PAR-6 at the tip of the sensilla cluster, indicating that it is the apical surface of these cells, which accumulates γ-tubulin::GFP similar to the pharynx and gut (Fig. 8 G, Fig. S6 G-L). We also observe α-catenin (HMP-1::GFP) at the furrow and midbody before it accumulates in the apical cluster of the sensilla neurons and remains there during dendrite extension (Fig. 8 H-J). To our knowledge, this is the first detailed examination of the division and initial steps of organization of these neuronal cell precursors.

**Figure 8.**
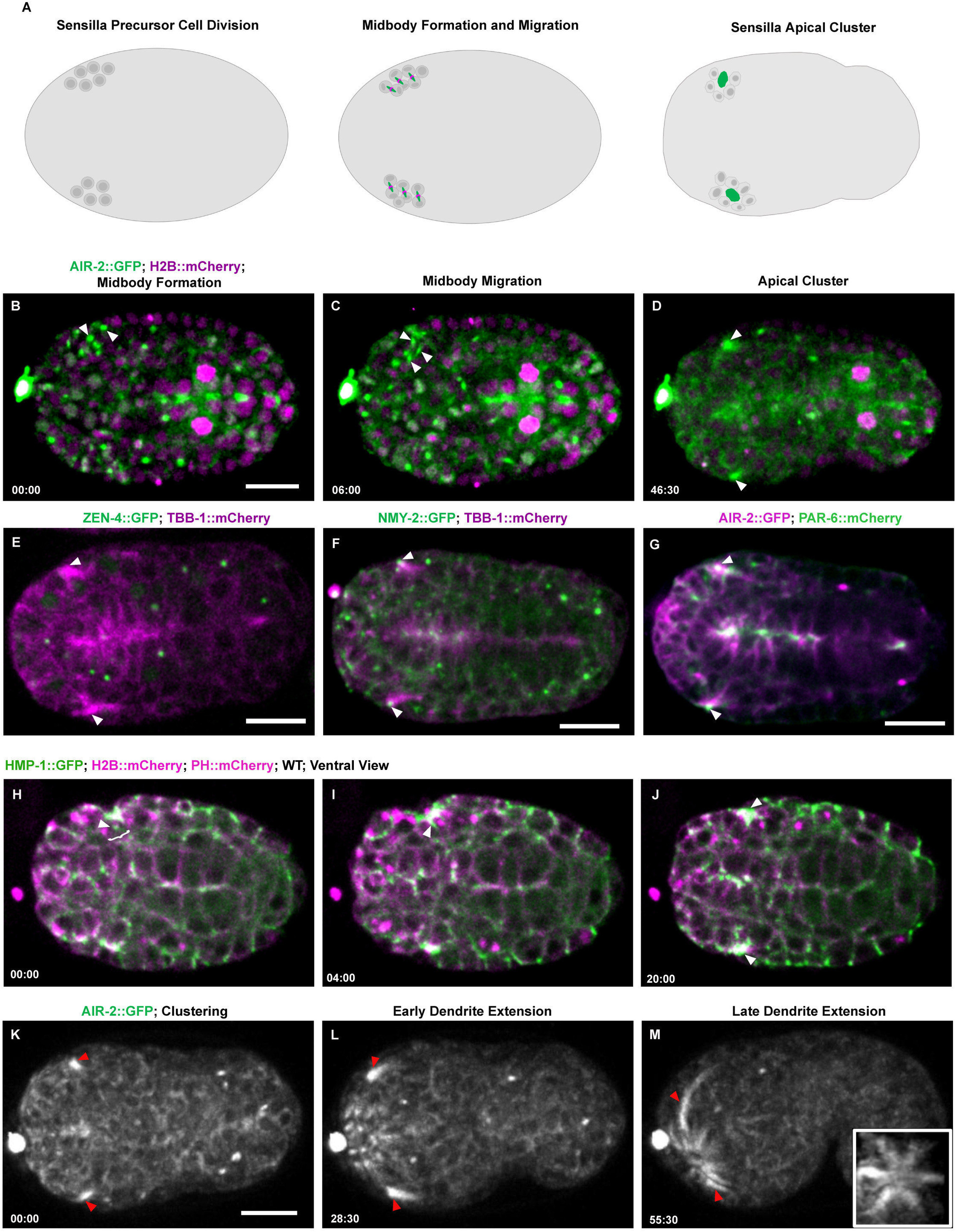
Midbody components label dendrites of sensilla neurons. (A) Diagram of sensilla precursor cell (SPC) divisions and the localization of the AIR-2::GFP at the midbody during cytokinesis until the apical clustering during polarization. (B-D) Cytokinesis in SPCs in the anterior lateral region of the embryo expressing AIR-2::GFP (green, H2B::Cherry in magenta) gives rise to multiple midbodies (arrowheads B,C) that cluster together at the lateral sides of the embryos (D). (E) The midbody ring marker ZEN-4::GFP (green, microtubules in magenta) is internalized and degraded before the cluster forms, which is also concentrated with microtubules (arrowheads). (F) NMY-2::GFP (green, microtubules in magenta) localizes to midbodies that cluster and remains at the tip of the dendrite as it extends. (G) PAR-6::mCherry (green, AIR-2::GFP in magenta) localizes to tip of the cluster (arrowheads) and persists at the tip of the dendrites as they extend, indicating that this is the apical surface of these cells. (H-J) HMP-1::GFP accumulates at the furrow and midbody (arrowhead, H, I) during sensilla precursor divisions and accumulates at the cluster and remains at the tip during dendrite extension (J). (K-M) After the apical cluster forms, AIR-2::GFP remains at the tip (red arrowheads) as the cells migrate to the nose of the animal. AIR-2::GFP also labels a substantial portion of the length of the dendrite as they extend (M). Insert in (M) is a rotated max z-projection showing the anterior end of the animal after multiple sensilla form. Time shown in minutes:seconds. Scale bar, 10 μm.

After formation of the cluster, we observe this apical region move and extend anteriorly until it reaches the tip of the animal during the bean stage through the late comma stage. AIR-2 remains localized along a substantial increasing length of the dendritic extension during the entire elongation process, as does tubulin (Fig. 8 K-M, Fig. S7 J-L, M-O and Video S12). As the amphid dendrites extend from the lateral sides of the embryo, other foci of AIR-2 form within the anterior region of the embryo and migrate toward the tip until six sensilla appear at the anterior tip (Fig. 8 M inset, and Video S12). Although the individual cell divisions cannot be easily discerned in this crowded anterior region, these data suggest that the sensilla in the tip of the animal form through a similar process. These results demonstrate that directly after cytokinesis a midbody migration event brings several midbody components to the apical tip of the amphid dendrites, which remain localized there as dendrite extension occurs. Neuronal cell polarization has been suggested to share mechanisms with epithelial morphogenesis (McLachlan and Heiman, 2013), suggesting that these modified cytokinesis events may play a role in cells that undergo epithelial polarization. Therefore, the midbody migrates from its original position at the end of furrowing to the apical surface in several developing tissues during morphogenesis. Interestingly, Aurora B remains localized at the apical surface of these tissues well after cytokinesis has occurred.

### Aurora B is required for proper dendrite formation

Given our observation cytokinesis in the developing sensory neurons we tested whether Aurora B kinase and other cytokinesis components were required for their formation. *C. elegans* amphid neurons can take up lipophilic dyes such as DiI when they form properly and generate cilia that are exposed to the environment (Hedgecock and White, 1985; Perkins et al., 1986). We maintained embryos at the permissive temperature (15 °C) and shifted them to the non-permissive temperature at different embryo stages until they hatched, then we stained the surviving L1 larvae with DiI. In wild-type, amphid neuron cell bodies, amphid dendrites, and phasmid neurons were clearly labeled by DiI and appeared normal as expected (Fig. 9 A). In the *air-2(or207)* mutant, we observed numerous defects in the subset of surviving embryos that did not fail to hatch and became L1 larvae (Fig. 9 B-E). Animals with no observed DiI staining were more common under longer inactivating conditions in the *air-2(or207)* mutant (Table 2). All *zen-4(or153)* fail to hatch when shifted during E4-E8, preventing analysis of DiI staining (Table 2). When shifted from the E8 stage, the few surviving *zen-4(or153)* larvae show severe DiI staining defects, which was dramatically reduced if embryos were shifted after the final divisions at the comma-1.5 fold stage (Fig. 9 F, Table 2). The *spd-1(oj5)* animals still had weak defects revealed by DiI staining despite having minimal cytokinesis failures, but never showed a complete lack of staining (Fig. 9 I, Table 2). These data are consistent with the hypothesis that proper execution of cytokinesis contributes to proper neurite development. Therefore, cytokinesis and AIR-2 function especially are required late in embryo development for epithelial polarization and morphogenesis of the lumen of the gut and proper formation of the sensilla neurons.

**Table 2.** Quantification of DiI Staining of Temperature-Sensitive Mutants

| Stage Before Shifting | Genotype | No DiI Signal | Weak DiI signal | Shape & Position Defect | Extended DiI Staining |
| --- | --- | --- | --- | --- | --- |
| 15 °C Forever | N2 | 0% (0/32) | 0% (0/32) | 0% (0/32) | 0% (0/32) |
|  | *air-2(or207)* | 2.7% (1/37) | 8.1% (3/37) | 2.7% (1/37) | 0% (0/37) |
|  | *zen-4(or153)* | 0% (0/18) | 0% (0/18) | 0% (0/18) | 0% (0/18) |
|  | *spd-1(oj5)* | 0% (0/34) | 0% (0/34) | 0% (0/34) | 0% (0/34) |
| E4-E8 | N2 | 0% (0/10) | 0% (0/10) | 0% (0/10) | 0% (0/10) |
|  | *air-2(or207)* | 50% (1/2) | 0% (0/2) | 50% (1/2) | 0% (0/2) |
|  | *zen-4(or153)* | 0% (0/0) | 0% (0/0) | 0% (0/0) | 0% (0/0) |
|  | *spd-1(oj5)* | 0% (0/44) | 4.5% (2/44) | 9.1% (4/44) | 6.8% (3/44) |
| E8-E16 | N2 | 0% (0/29) | 0% (0/29) | 0% (0/29) | 0% (0/29) |
|  | *air-2(or207)* | 16.7% (2/12) | 58.3% (7/12) | 58.3% (7/12) | 0% (0/12) |
|  | *zen-4(or153)* | 77.8% (7/9) | 11.1% (1/9) | 22.2% (2/9) | 0% (0/9) |
|  | *spd-1(oj5)* | 0% (0/59) | 0% (0/59) | 8.5% (5/59) | 5.1% (3/59) |
| Comma-1.5 Fold | N2 | 0% (0/21) | 0% (0/21) | 0% (0/21) | 0% (0/21) |
|  | *air-2(or207)* | 6.9% (2/29) | 10.3% (3/29) | 0% (0/29) | 13.8% (4/29) |
|  | *zen-4(or153)* | 0% (0/53) | 0% (0/53) | 0% (0/53) | 22.6% (12/53) |
|  | *spd-1(oj5)* | 0% (0/18) | 0% (0/18) | 0% (0/18) | 33.3% (6/18) |

**Table 3.**
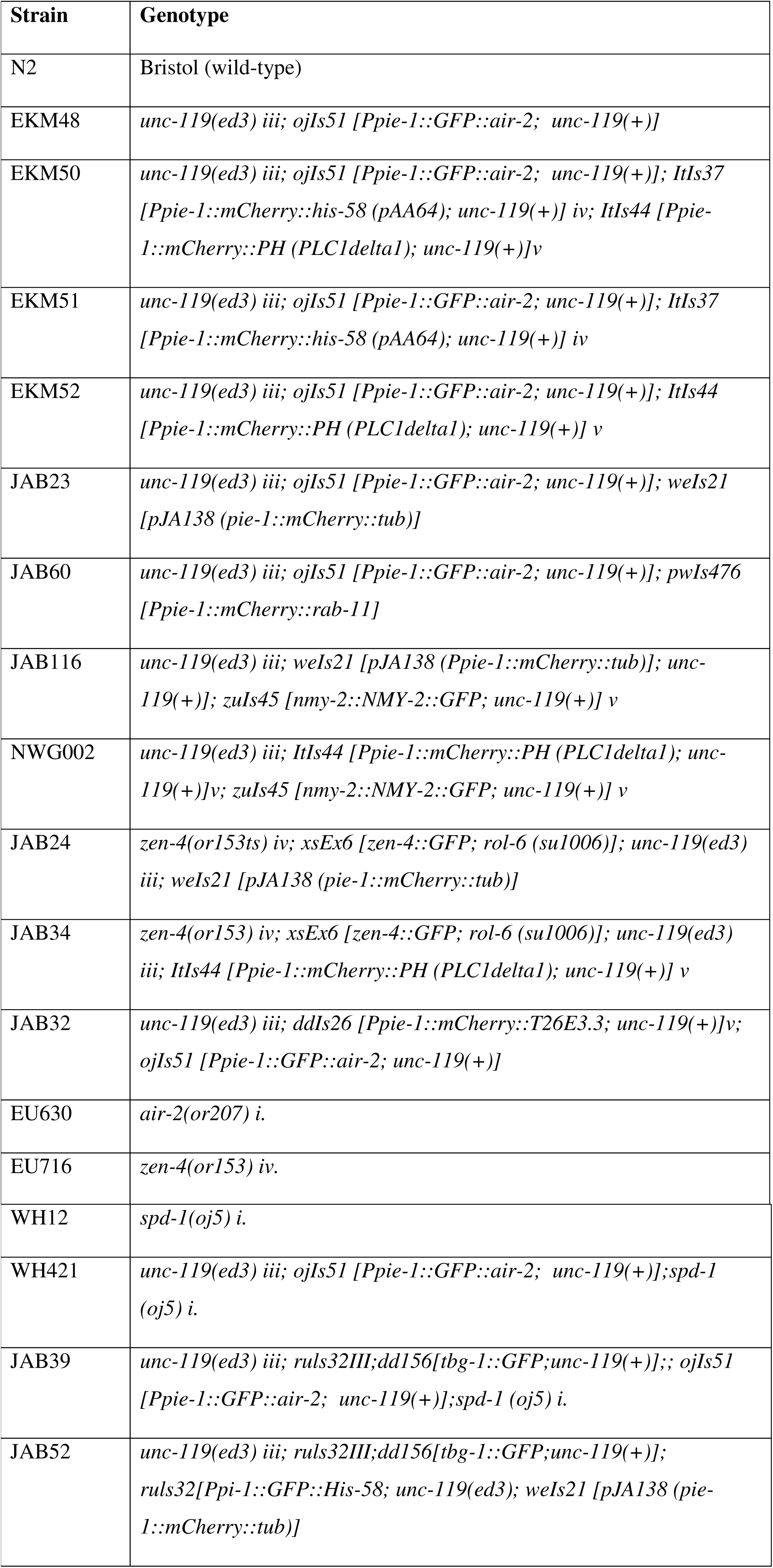
Strains used in this study.

**Figure 9.**
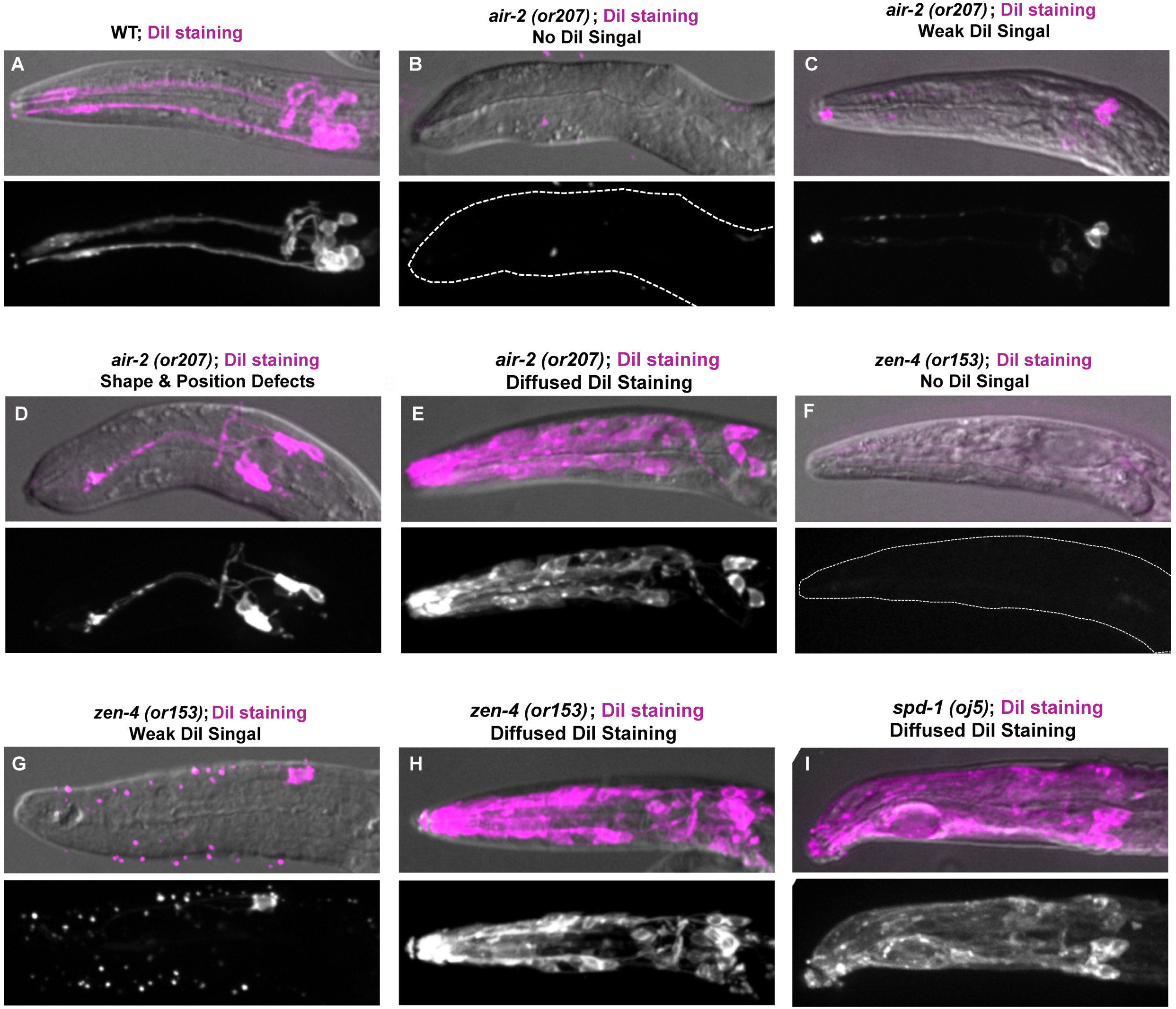
Cytokinesis mutants have disrupted sensilla neuron morphology. Visualizing dendrite and neuron morphology by DiI staining in surviving larvae after shifting mutants to 25 °C at the E4 or E8 stage until hatching. (A) In wild-type animals, two dendrite bundles can be clearly observed as well as amphid and phasmid neurons. (B-E) Hatched mutant larvae displayed a variety of neurite defects, including No-DiI signal (B, F), Weak signal (C, G), dendrite shape and positioning defects (D) and additional diffuse staining throughout the head of the animal (E, H, I).

## Discussion

Our results have revealed complex and reproducible patterns of cytokinesis during the invariant embryonic divisions in *C. elegans.* The entire invariant lineage has been known for several decades and our results suggest that cytokinesis also follows a specific pattern during the lineage. We observe reproducible alterations to furrow symmetry, central spindle length, abscission timing, midbody movement and inheritance. The traditional view of the embryo lineage is that cells are born and subsequently undergo changes that produce the differentiated organization within a tissue. However, our data demonstrate that cells in multiple tissues are completing cytokinesis when they polarize during morphogenesis. We also demonstrate that failures of cytokinesis disrupt proper epithelial polarization, disrupting nuclear positioning and apical accumulation of microtubules and adhesion complexes. This role for cytokinesis during polarization might explain why many cells in the lineage, including several of the amphid neuronal precursors, divide and produce one daughter cell that undergoes apoptosis instead of finishing the divisions earlier when the right number of cells are generated. Such a role for a modified cytokinesis is observed later in the Q neuroblast that generates a smaller a daughter cell that undergoes apoptosis, which is prevented if the parameters of cytokinesis change (Ou et al., 2010). Given that the entire cell is reconfigured during mitosis and that cytokinesis is the transition period back into the interphase state, this is an ideal time window to reorganize cellular architecture. Understanding how the developmental plasticity of cytokinesis is regulated will be a fascinating question for future studies.

We observe consistent changes to the symmetry of furrow ingression where the first mitosis is relatively symmetric and the second mitosis is highly asymmetric. Previously, the furrow asymmetry in the first division was shown to be a consequence of asymmetric accumulation of contractile ring components during ingression (Maddox et al., 2007). The adhesion between cells may also reinforce this asymmetry to drive the highly asymmetric furrow observed in the second round of divisions (Padmanabhan and Zaidel-Bar, 2017). The dynamic localization of adhesion we observe during cytokinesis at several stages of development suggest that adhesion may control cellular interactions during the disruptive process of anaphase spindle elongation and cytokinesis to maintain cellular positioning in the embryo. Whether due to cell intrinsic or extrinsic factors, the asymmetric furrows have previously been postulated to drive efficient furrowing or help maintain proper cell-cell contacts during cytokinesis (Maddox et al., 2007; Morais-de-Sa and Sunkel, 2013). Our data suggest another hypothesis: the asymmetric furrow may be required for the AB midbody to be engulfed by EMS instead of either daughter cell. Given that the midbody has been proposed to deliver signals to cells that inherit it, it is worth noting that the MS cell collects up to four midbodies over time (Singh and Pohl, 2014). We see relatively symmetric furrowing in several tissues later in morphogenesis. This is unexpected because an asymmetric furrow would be sufficient to position the midbody at the nascent apical surface. Given that the polarization mechanisms are not completely understood, for example the extracellular matrix component laminin is required in the pharynx but not the intestine (Rasmussen et al., 2012), the symmetrical furrow followed by midbody migration may be important for defining and positioning the apical surface. Perhaps there is no good reference for an asymmetric furrow to position the midbody at the apical surface prior to epithelial polarization in cells in different locations. We hypothesize that lumen formation in the gut and pharynx is analogous to that described in MDCK cells with the formation of a midbody-derived apical-membrane initiation site with the addition of midbody migration for correct positioning of this domain (Li et al., 2014).

The coordinated, directed movement of the midbody we observe in several tissues represents a new phenomenon during cytokinesis. Our data also suggest that abscission has not taken place before the midbody migrates in the intestine. This would mean that the two daughter cells polarize while connected at the midbody, which might facilitate their reorganization. These data are somewhat different from what is observed in already polarized epithelia where the furrow constricts from the basal to the apical surface to position the midbody. We demonstrate that Aurora B is required for successful completion of cytokinesis and polarization of microtubules and adhesion. In the future it will be important to specifically prevent the movement of the midbody and determine the affect it has on the polarization process. We hypothesize that midbody migration to the apical surface is involved in cells that polarize during a mesenchyme to epithelial transition. Previously, midbodies have been shown to reposition after forming under normal or mutant conditions (Bernabe-Rubio et al., 2016; Herszterg et al., 2013; Morais-de-Sa and Sunkel, 2013; Singh and Pohl, 2014), but this phenomena is only appreciated in isolated cases and poorly understood. The entire cortex is controlled by several actin cytoskeletal regulators in order to perform cytokinesis (Jordan and Canman, 2012), perhaps this is also employed to control the movement of the midbody. In the future, it will be important to investigate how the midbody moves to the apical surface and how various delivered proteins contribute to polarization.

In the tissues we investigated, the cells are undergoing their terminal cell division before morphogenesis, although some cells like those in the gut undergo post-embryonic divisions. These cells are also undergoing epithelial polarization and a mesenchymal to epithelial transition. After midbody movement, RAB-11, AIR-2, microtubules and possibly other molecules are recruited to the apical surface. Certainly, these different tissues have unique gene expression programs, part of which might involve proteins delivered to the midbody and the apical surface. For example, a transmembrane protein that binds to an extracellular partner is expressed in the tip of the dendrites in amphid sensilla, which is required to maintain dendrite attachment at the tip of the embryo (Heiman and Shaham, 2009). It is unknown how this protein localizes to the tip of the dendrite, but one speculative possibility given our observations that midbody components localize to the same site is that it could be delivered through cytokinesis-directed membrane trafficking. Delivering proteins through the midbody was observed in neuroepithelial cells where a stem cell marker protein is released into the extracellular environment in released midbody remnants (Dubreuil et al., 2007). Later in life, the worm releases exosomes from the sensory cilia that form at the tip of the dendrites of the sensilla for communication between animals (Wang et al., 2014a). Perhaps the initial secretory apparatus built during cytokinesis to promote cell division is recruited to the apical surface of these neurons to enable exosome release. Further investigation is required to define the molecular contributions provided by the midbody to the apical surface of these tissues.

Once the midbody moves to the apical surface, we observe that different components of the midbody have different fates, which is an unexpected and novel observation. Typically, once the midbody is abscised from the cell, it is thought that most midbody proteins are discarded with the remnant, as observed in the early embryo divisions. Aurora B kinase remains at the apical surface well after other midbody components like ZEN-4 are removed. The limit of the resolution of light microscopy does not allow us to characterize in detail how this occurs. The most likely model is that the central midbody bulge is cut from the plasma membrane and flank proteins like Aurora B, RAB-11, and microtubules are left behind. Among the many mitotic functions of Aurora B, it is a critical regulator of the timing of abscission (Mathieu et al., 2013; Steigemann et al., 2009). Based on our observations of midbody flank microtubules and the timing of midbody ring internalization, abscission likely occurs after the midbody migration event, and the delay in abscission might require Aurora B activity. Delayed abscission by Aurora B kinase in mouse embryos was shown to maintain interphase bridges that mediate RAB-11 dependent delivery of cell adhesion molecules to apical membranes (Zenker et al., 2017). We find that inhibition of Aurora B diminishes spindle midzone microtubules and leads to reduced adhesion at the furrow and midbody during cytokinesis, which is consistent with such a mechanism operating during epithelial polarization. Aurora B was recently shown to be regulated by an atypical cadherin in early zebrafish embryos to regulate the spindle midzone microtubule organization (Chen et al., 2018), further suggesting reciprocal regulation between adhesion and cytokinesis. Aurora B also regulates a number of cytoskeletal regulators during cytokinesis that control cell shape (Ferreira et al., 2013; Floyd et al., 2013; Goto et al., 2003; Kettenbach et al., 2011), and it will be important to determine whether any are involved with the events we observed. In the intestine, the central spindle elongates dramatically as the midbody migrates, which might also be regulated by Aurora B (Bastos et al., 2013). Spindle midzone regulation is controlled by altered expression of the central spindle protein PRC-1 (the homologue of *spd-1*) in the *Xenopus* embryo, which correlates with changes to furrow ingression and midbody behavior (Kieserman et al., 2008). While we observe the centralspindlin component ZEN-4 becoming rapidly internalized with the midbody in the three tissues, it was previously implicated in morphogenesis of the epidermis and pharynx (Hardin et al., 2008; Portereiko et al., 2004; Von Stetina et al., 2017). It remains to be determined whether this role is related to the dynamics of cytokinesis or a cytokinesis-independent function of ZEN-4 as previously suggested. Therefore, further study will be required to understand the role of the spindle midzone components during the specialized cytokinesis events that occur during morphogenesis.

In the sensilla, the centriole moves to the tip of the dendrite to form the base of the sensory cilia of these neurons (Dammermann et al., 2009; Nechipurenko et al., 2017; Perkins et al., 1986). Multiple central spindle proteins localize to the base of cilia in *Xenopus* epithelial cells and are required for cilia morphology after the divisions are completed in *C. elegans* (Kieserman et al., 2008; Smith et al., 2011). Additionally, loss of Aurora B kinase causes aberrant neuronal axon morphology, and overexpression of Aurora B causes extended axonal outgrowth in zebrafish (Gwee et al., 2018). At the apical surface of the gut, γ-tubulin and other pericentriolar material is delivered from the centrosome while the centrioles are discarded (Feldman and Priess, 2012). The gut apical membrane ultimately becomes elaborated with microvilli (Feldman and Priess, 2012; Leung et al., 1999). We also observed γ-tubulin at the apical surface of the pharynx and sensilla dendrites. Therefore, different material provided by the midbody and centrosome may contribute to the cytoskeletal architecture of the apical surface depending on the needs of each tissue. Delineating the precise relationship between these two organelles and deciphering how cytokinesis contributes to proper cellular reorganization during morphogenesis will be a major focus of future studies.

## Acknowledgements

Lattice light sheet microscopy was performed in collaboration with the Advanced Imaging Center at HHMI Janelia Research Campus, a facility jointly supported by the Gordon and Betty Moore Foundation and the Howard Hughes Medical Institute. We appreciate the CGC and Wormbase funded by the NIH Office of Research Infrastructure Programs (P40 OD010440) and National Human Genome Research Institute (U41 HG002223), which provided some *C. elegans* strains and genome information. We are grateful to members of the Bembenek laboratory for productive discussion, reagent preparation and handling strains. We also thank Dr. Max Heiman, Dr. Zhirong Bao for discussions and Dr. Don Fox, Dr. John White, Dr. John Heddleston, Dr. Heidi Hehnley-Chang, Lindsay Rathbun, and Erica Colicino for critical feedback on the manuscript. Funding was provided by the Ministry of Science Technology of Taiwan (105-2119-M-001-026-MY2) to C.B.C., the Howard Hughes Medical Institute to E.B., and NIH R01 GM114471 to J.N.B.

## Materials and Methods

### *C. elegans* Strains

*C. elegans* strains were maintained with standard protocols. *C. elegans* strains expressing midbody proteins driven by the *pie-1* promoter are listed in Table 2. All temperature-sensitive mutants were obtained from the Caenorhabditis Genetics Center.

### Embryo Preparation and Imaging

For live imaging, young gravid hermaphrodites were dissected in M9 buffer containing polystyrene microspheres and sealed between two coverslips with vaseline (Pohl and Bao, 2010). Live cell imaging was performed on a spinning disk confocal system that uses a Nikon Eclipse inverted microscope with a 60 X 1.40NA objective, a CSU-22 spinning disc system, and a Photometrics EM-CCD camera from Visitech International. Images were acquired by Metamorph (Molecular Devices) and analyzed by ImageJ/FIJI Bio-Formats plugins (National Institutes of Health) (Linkert et al., 2010; Schindelin et al., 2012). Whole embryo live imaging was performed on a lattice light sheet microscopes housed in the Eric Betzig lab, Bi-Chang Chen lab, or the Advanced Imaging Center at HHMI Janelia. The system is configured and operated as previously described (Chen et al., 2014). Briefly, embryos were dissected out and adhered to 5 mm round glass coverslips (Warner Instruments, Catalog # CS-5R). Samples were illuminated by lattice light-sheet using 488 nm or 560 nm diode lasers (MPB Communications) through an excitation objective (Special Optics, 0.65 NA, 3.74-mm WD). Fluorescent emission was collected by detection objective (Nikon, CFI Apo LWD 25XW, 1.1 NA), and detected by a sCMOS camera (Hamamatsu Orca Flash 4.0 v2). Acquired data were deskewed as previously described (Chen et al., 2014) and deconvolved using an iterative Richardson-Lucy algorithm. Point-spread functions for deconvolution were experimentally measured using 200nm tetraspeck beads adhered to 5 mm glass coverslips (Invitrogen, Catalog # T7280) for each excitation wavelength.

### Immunostaining Assay in *C. elegans* Embryos

Apical marker staining was performed with the freeze-crack methanol protocol (Leung et al., 1999). Immunostaining with anti-AIR-2 antibodies was performed as described (Schumacher et al., 1998). Primary antibodies and (dilutions) used were anti-ERM-1 (1:200); P4A1/PAR-3 (1:200); DLG-1 (1:200); MH33 (1:150); AIR-2 (1:50). 1:200-400 dilutions of Alexa 588 and 468 secondary antibodies were used in the study. To stain temperature-sensitive mutants, two-cell stage embryos were dissected from gravid worms, mounted in 10 μL of M9 buffer, and kept cold on ice. The two-cell stage embryos were incubated at 15 °C for 4-7 hours until specific stages, then shifted to the restrictive temperature (25 °C) for 2-4 hours and stained as described above.

### DiI staining in *C. elegans*

DiI staining of wild-type and temperature sensitive mutants was done as previously described (Tong and Burglin, 2010). Two-cell stage embryos were incubated at 15 °C for 6.5∼7 hours until they reached the polarized E16 stage, then shifted to the restrictive temperature (25 °C) with 1:200 dilution of stock DiI dye solution containing 2 mg/mL DiI in dimethyl formamide for 18-24 hours. Hatched larvae were transferred to M9 and washed twice in M9 before mounting in 25 mM levamisole on 2% agar pads for imaging.

### Temperature-Shift Experiments

Temperature-sensitive mutants were maintained at 15 °C. To perform temperature shifts on staged embryos, gravid adults were transferred to a dissection chamber (< 4 °C), which was precooled in ice bucket, with 20 μL of ice-cold M9 Buffer. Two-Cell stage embryos were quickly transferred (within a 5-10 minute time window) via mouth pipette (Aspirator tube assemblies, Sigma) to Fisherbrand Hanging Drop Slides (Catalogue #12-560B) on ice. The slide was placed into a humidified chamber and incubated at 15 °C until the appropriate stages were reached and then shifted to 26 °C. Incubation times were determined based on *C. elegans* embryonic lineage timing and adjusted according to DAPI staining to ensure each mutant was shifted at a similar stage of embryo development. To inactivate *air-2(or207)*, mutant embryos were incubated for 5 hours at 15 °C and shifted to 26 °C for 3 hours to reach the bean stage or 5 hours at 26 °C to reach the comma stage. This was the minimum amount of time to shift embryos to non-permissive temperature and observe significant cytokinesis defects by the E8-E16 division, indicating significant reduction of AIR-2 function. Most embryos reached the E4-E8 division at the time of the shift. By live imaging we found that there was little disruption of the E4-E8 division under these conditions since (n=4/5) *air-2(or207)* embryos have 8 normal E8 cells. N2, *spd-1(oj5)*, and *zen-4(or153)* embryos were incubated for 4.5 hours at 15 °C to reach E4-E8 stage, followed by 3 hours at 26 °C to reach the bean stage and 5 hours at 26 °C to reach the comma stage. To shift embryos at the comma stage, *air-2(or207)* embryos were incubated for 12 hours at 15 °C and N2, *spd-1 (oj5) and zen-4(or153)* embryos were incubated 11-11.5 hours at 15 °C.

## SUPPLEMENT FIGURE LEGENDS

**Figure S1.**
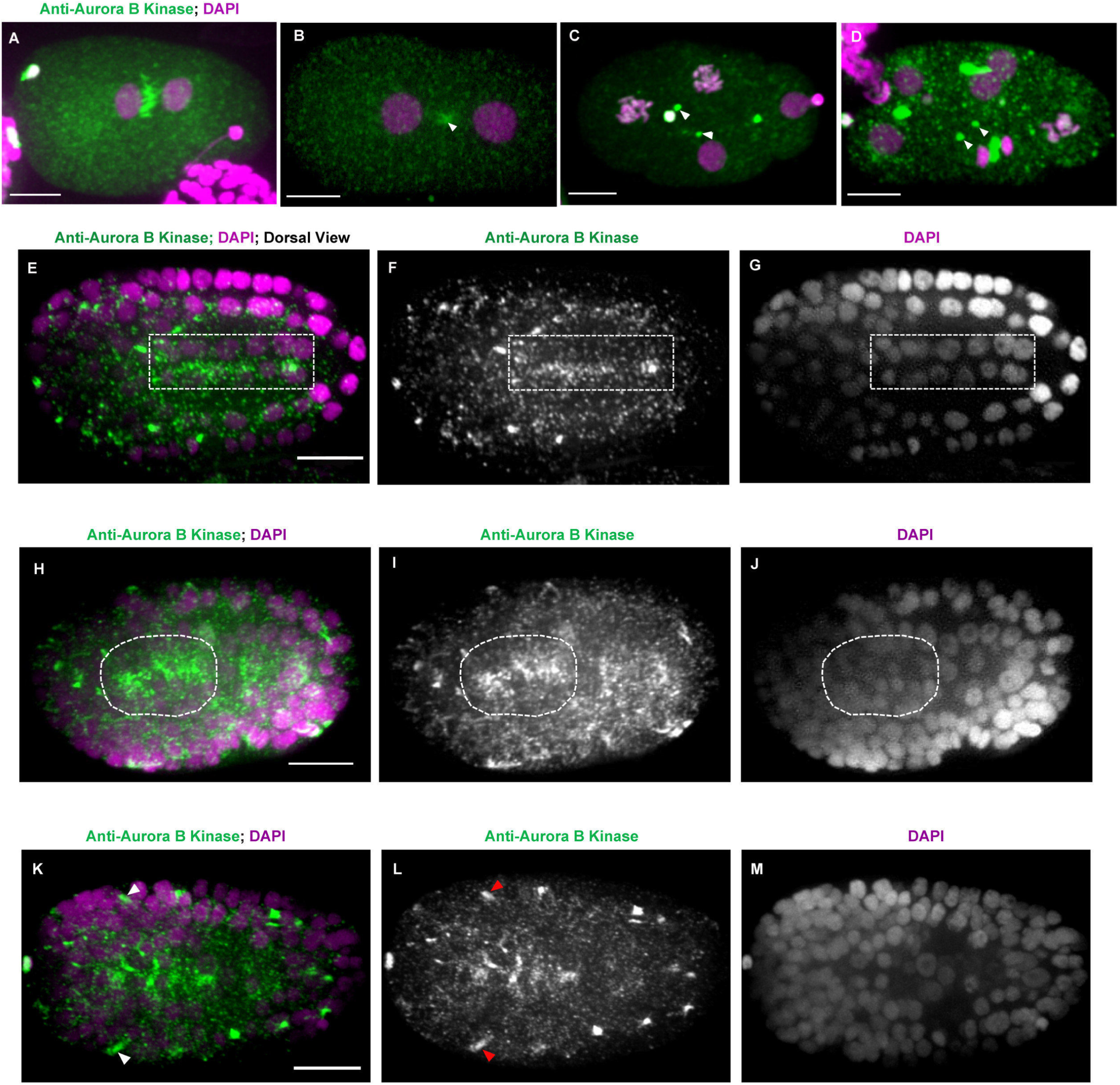
Immunostaining of endogenous Aurora B kinase during embryo development. Immunostaining of Aurora B kinase (green) with DAPI (magenta) in wild type embryos. In early embryos, AIR-2 is observed at the central spindle (A), midbody flank (arrowhead in B), and midbody remnants (arrowheads in C, D). (E-G) Polarized E16 wild type embryo shows localization at the apical surface of the intestine (E-G, rectangle). Staining of wild type bean stage embryos shows apical localization in the pharynx (H-J, dotted circle) and at the apical cluster of the amphid sensilla (K-M, arrowheads). Scale bar, 10 μm.

**Figure S2.**
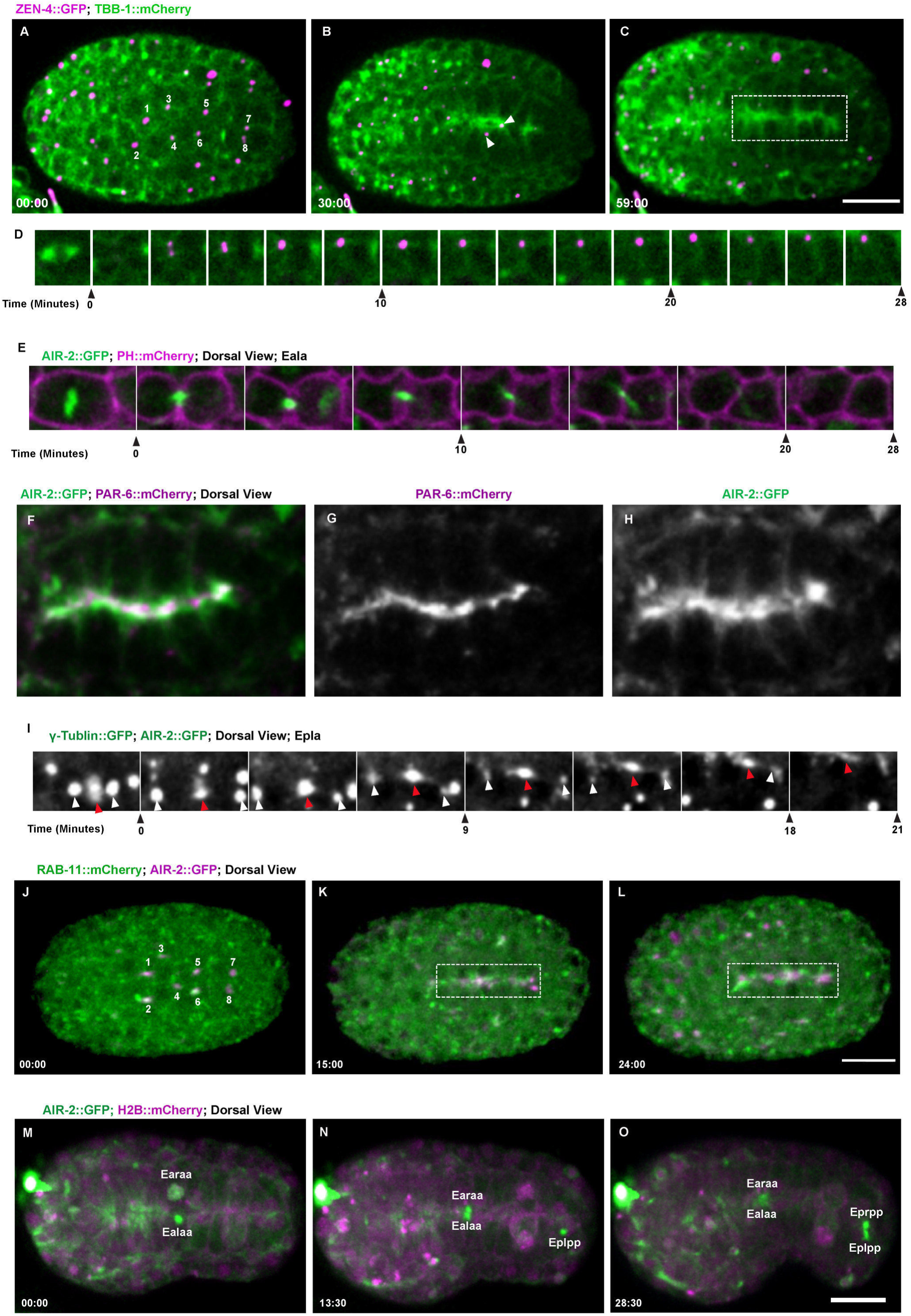
Cytokinesis in the intestine epithelia Cytokinesis in the E8-E16 division. (A-C) ZEN-4::GFP (magenta, TBB-1::mCherry in green) appears on midbodies (A, numbered 1-8) that migrate to the midline (B) and are quickly removed (C, rectangle box). (D) Single plane imaging tracking the movement of ZEN-4::GFP (magenta,TBB-1::mCherry in green) to the apical surface. (E) Midbody from Eala at the E8-E16 division migrates towards the midline but the AIR-2 signal diminishes (green, PH::mCherry in magenta). (F-H) The apical surface marker, PAR-6::mCherry (G, magenta in F), colocalizes with AIR-2::GFP (H, green in F) at the apical midline. (I) Image series showing the simultaneous movement of AIR-2::GFP (red arrowhead) on the midbody and y-tubulin::GFP (white arrowheads) on centrosomes to the apical surface. (J-L) RAB-11::mCherry (green) and AIR-2::GFP (magenta) colocalize on midbodies (labeled as 1-8 in J) as they migrate to the midline (K) and persist well after polarization is complete (L, rectangle). (M-O) In the terminal E16-E20 division, AIR-2::GFP appears on four midbodies (labeled according to E16 mother cell identity) that move to the apical midline (N, O). Time shown in minutes: seconds. Scale bar, 10 μm.

**Figure S3.**
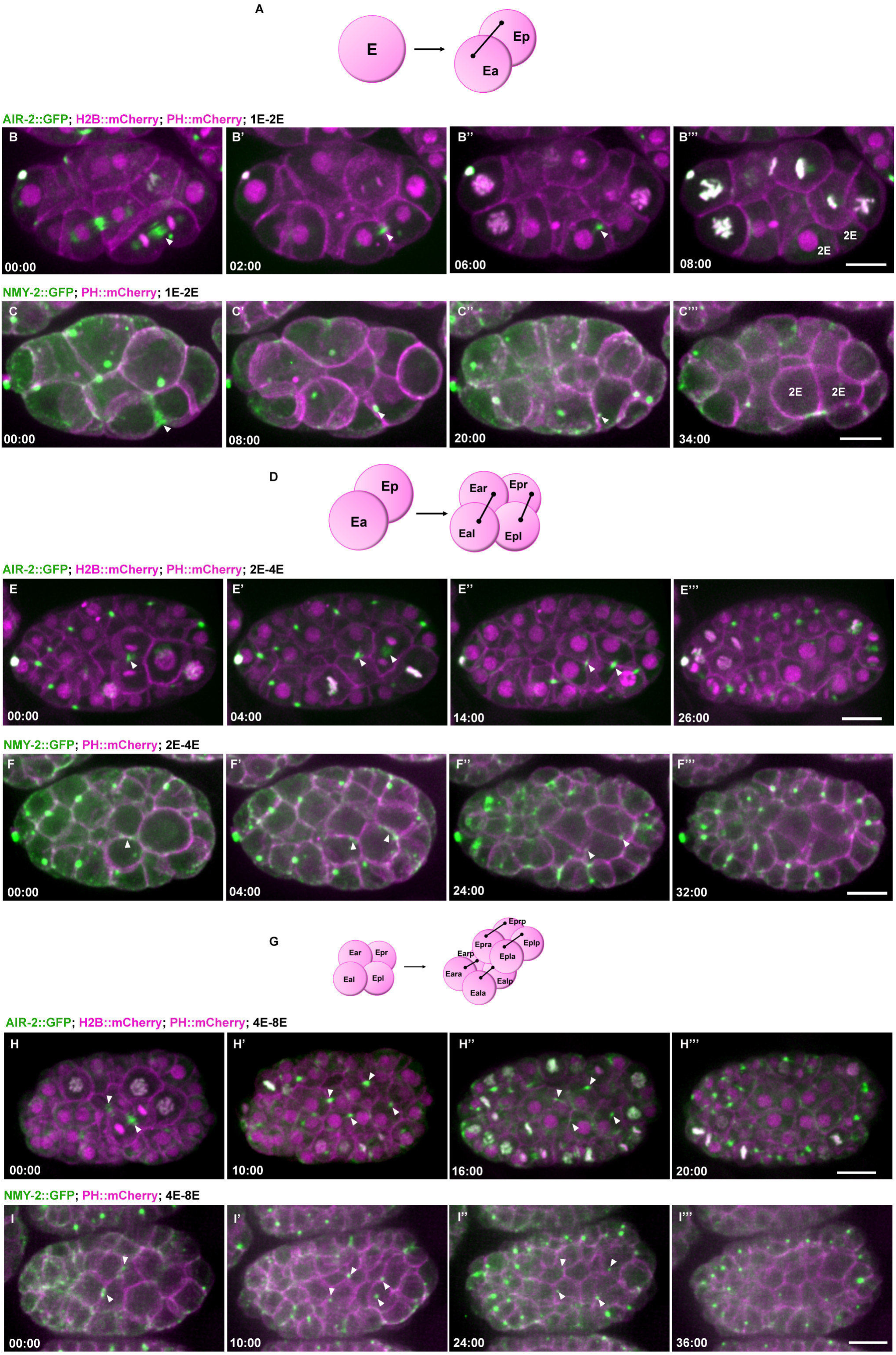
Midbody dynamics during divisions of the early E lineage. (A) Diagram of the E-E2 division. Dynamics of Aurora B (B) and NMY-2 (C) show formation of the midbody (arrowheads) and rapid internalization. A similar pattern is also observed during the E2-E4 divisions (D-F) and the E4-E8 divisions (G-I), demonstrating that the early embryo divisions of the E lineage are similar until morphogenesis and epithelial polarization. Scale bar, 10 μm.

**Figure S4.**
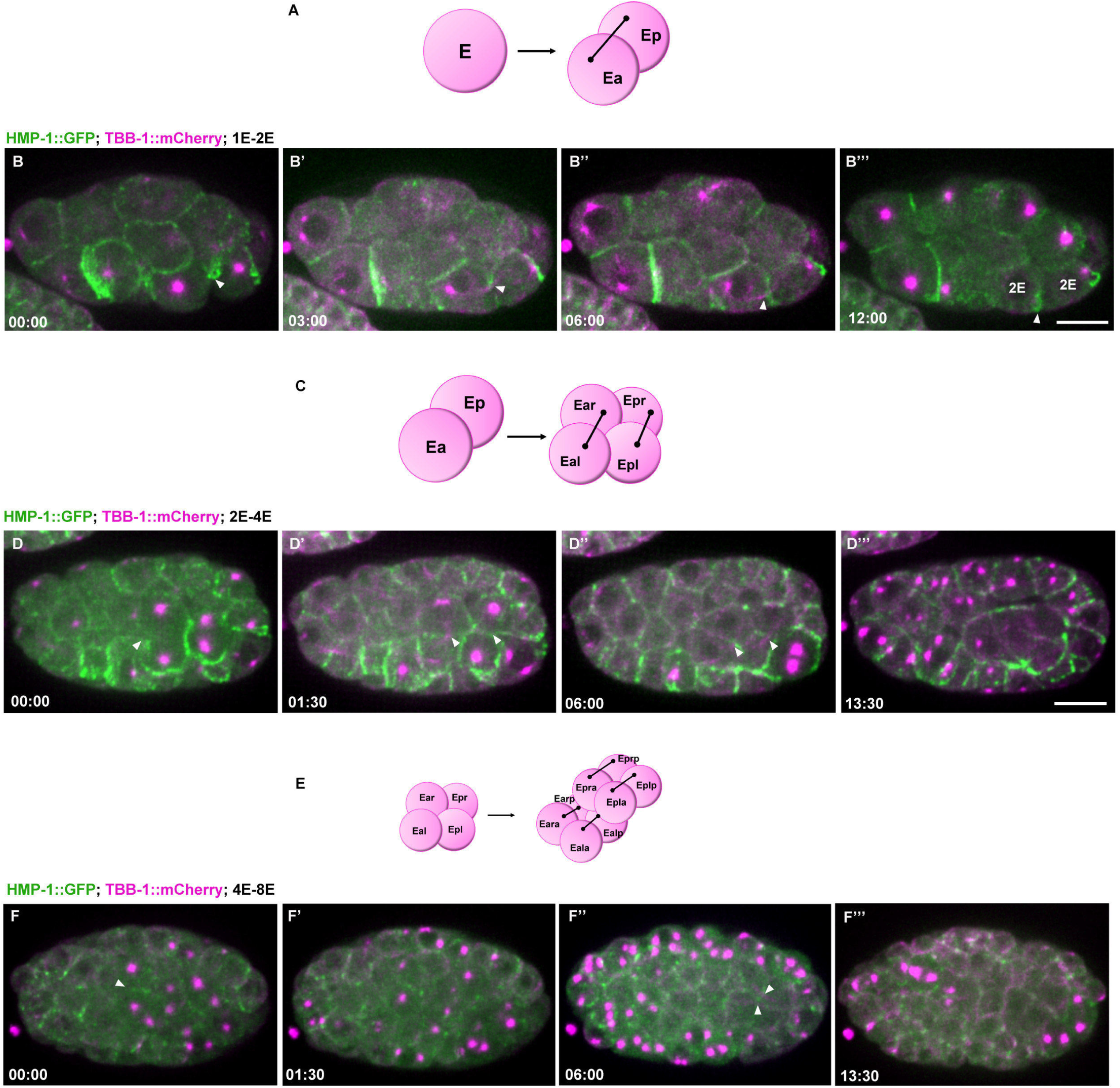
Furrow and midbody localization of α-catenin during cytokinesis in the E lineage divisions. (A) Diagram of the E-E2 division. HMP-1::GFP localizes to the furrow and membrane adjacent to the midbody during cytokinesis (arrowheads in B). Shortly after this, apical constriction promotes gastrulation of the gut cells into the core of the embryo. (C) Diagram of the E2-E4 division. HMP-1::GFP can be observed in the cortex of the dividing cells and localizes along the furrow membrane for a short time after cytokinesis (arrowheads, D). (E) Diagram of the E4-E8 divisions. (F) Arrowheads indicate accumulation of HMP-1::GFP near the midbody during cytokinesis. HMP-1::GFP does not accumulate at the midline until the end of the E8-E16 division. Scale bar, 10 μm.

**Figure S5.**
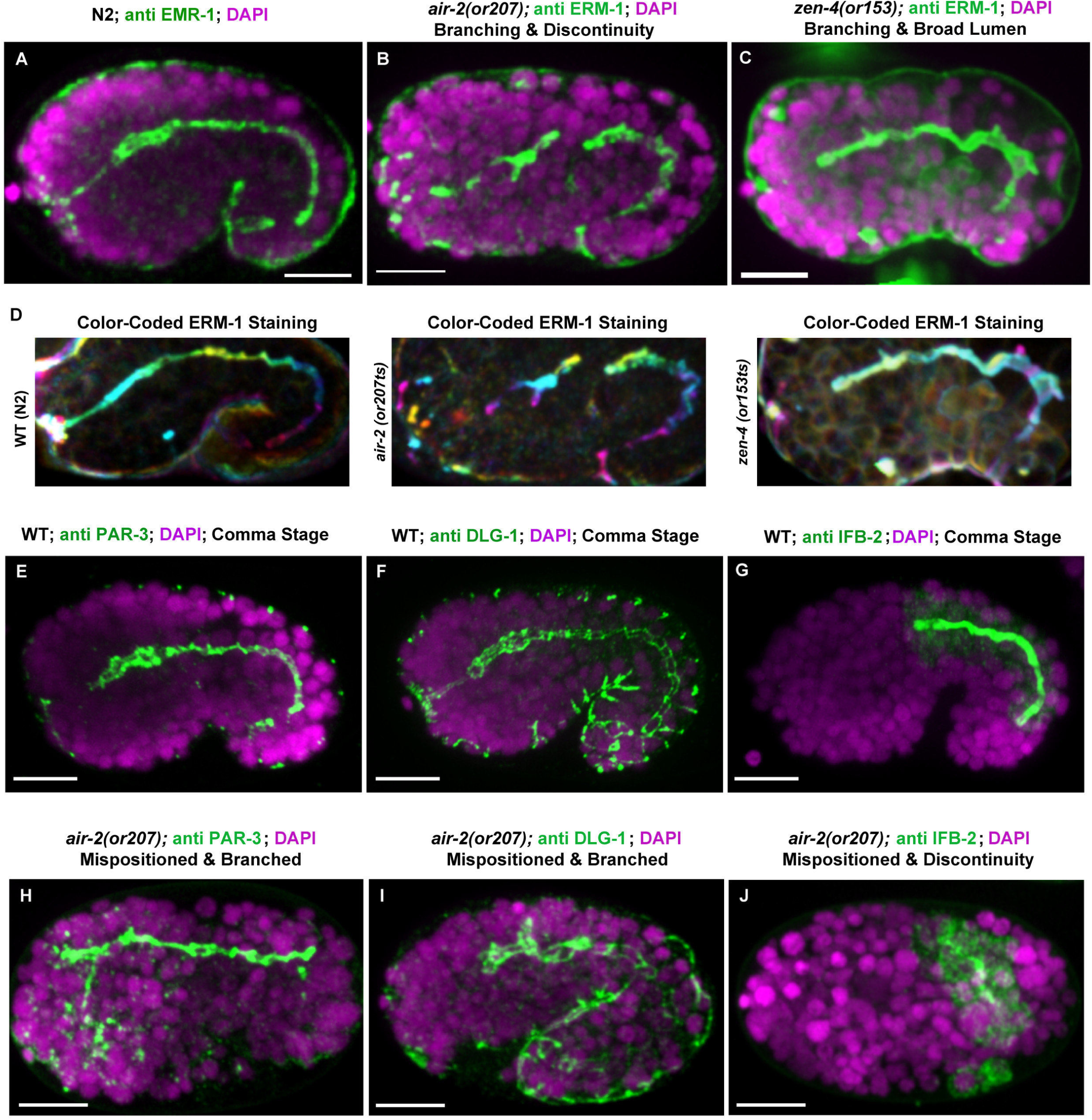
Cytokinesis mutants have disrupted intestinal and pharyngeal tubulogenesis. (A) ERM-1 stains the apical lumen of the gut and pharynx in embryos at the comma stage. Persistence of gut lumen defects are observed in (B) *air-2(or207)* and (C) *zen-4(or153)* embryos. (D) Color-coded max Z-projections of A-C. (E-G) Staining of multiple apical surface markers in WT embryos shows normal localization of (E) PAR-3, (F) DLG-1 and (G) IFB-2. In *air-2(or207)* mutants, apical surface markers localize to distorted and mispositioned lumens (H-J). Scale bar, 10 μm.

**Figure S6.**
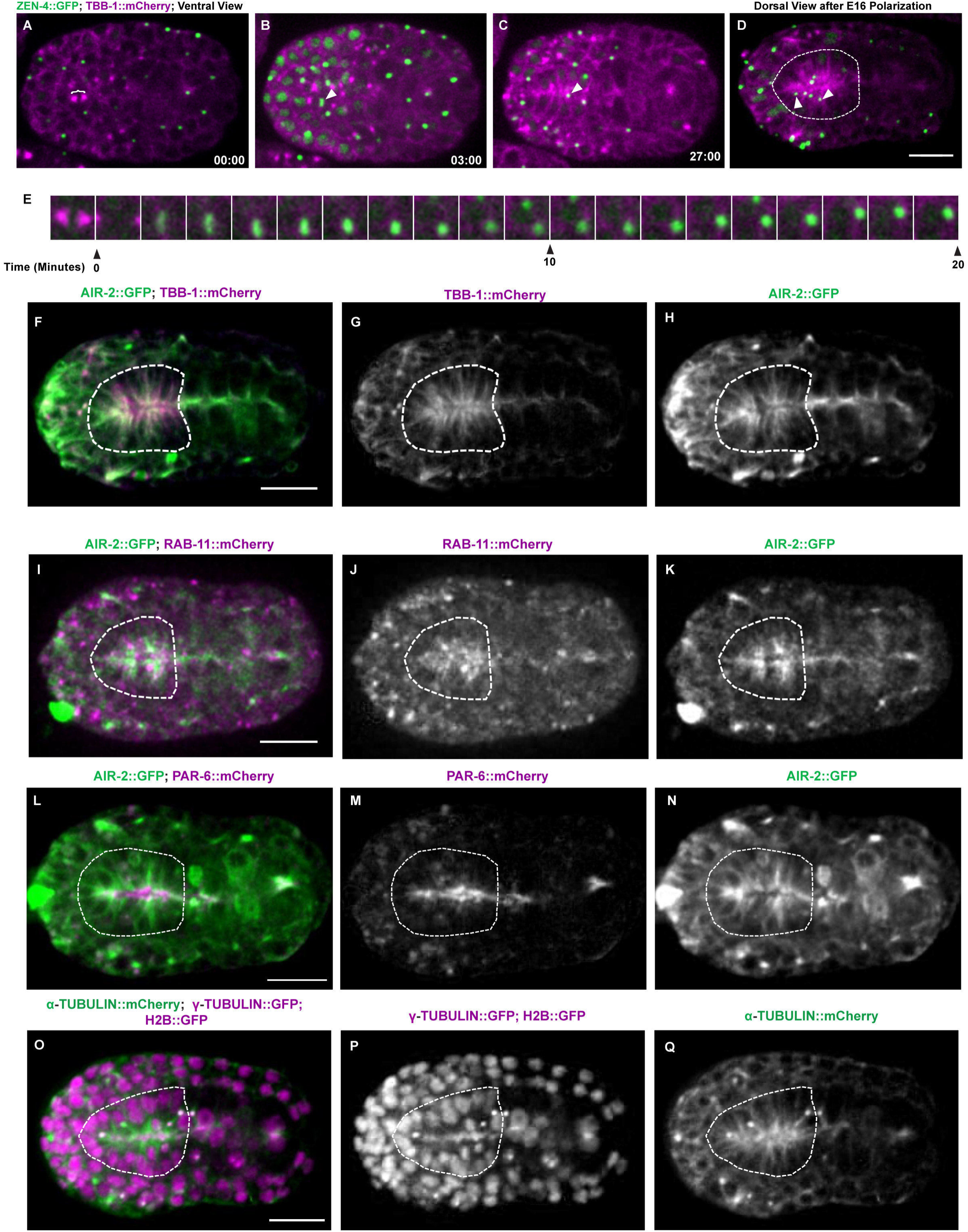
Cytokinesis in Pharynx Precursor Cells. (A-E) Centralspindlin ZEN-4::GFP dynamics in PPC divisions. Midbody remnants labeled with ZEN-4::GFP migrate to the midline (arrowheads, C) and are rapidly internalized and degraded. (F-K) AIR-2::GFP (H, K, green in merge) colocalized with TBB-1::mCherry (G, magenta in merge) and partially colocalized with RAB-11::mCherry (J, magenta in merge) at the apical surface of the polarized pharynx (dotted circle). (L-N) AIR-2::GFP (N, green in merge) partially co-localized with PAR-6::mCherry (M, magenta in merge) at the apical surface of the pharyngeal bulb (dotted circle). (O-Q) The apical surface of the pharynx accumulates γ-tubulin::GFP (P, magenta in merge, microtubules in green). Scale bar, 10 μm.

**Figure S7.**
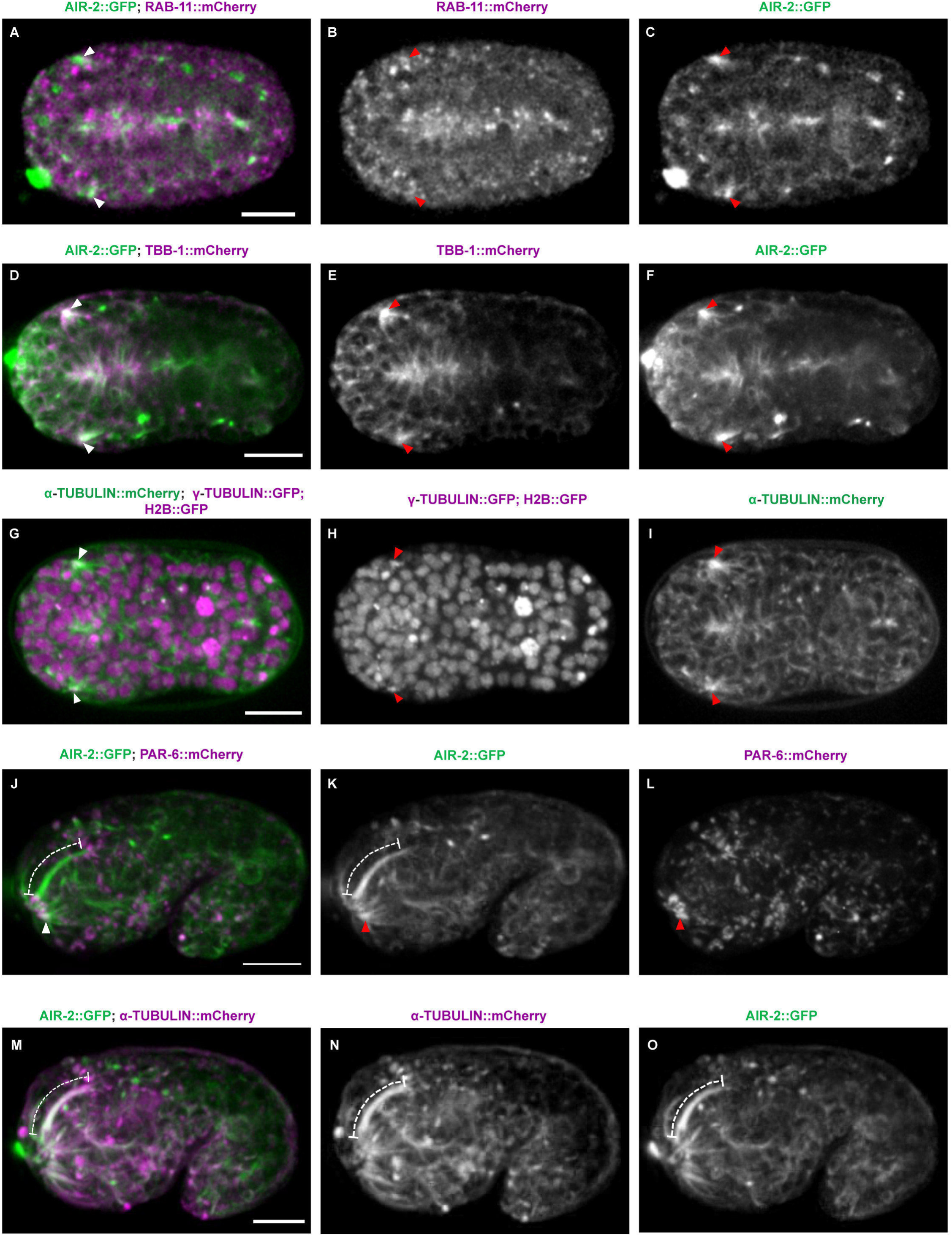
Midbody components label dendrites of sensilla neurons. Aurora B kinase (green) colocalizes with (A-C) RAB-11::mCherry (magenta) and (D-F) TBB-1::mCherry (magenta) at the apical cluster in forming sensilla neurons (red arrowheads). (G-I) γ-tubulin::GFP (H, magenta in merge) localizes at the apical cluster with microtubules (I, green in merge). (J-L) AIR-2::GFP (green) localizes along the dendrite extension (dashed line) and PAR-6::mCherry is observed at the tip of the extension (arrowhead), indicating it is the apical domain. (M-O) AIR-2::GFP (green) and TBA-1::mCherry (magenta) colocalize along the length of the extended dendrite (dashed line). Scale bar, 10 μm.

## SUPPLEMENT MOVIE LEGENDS

### Video S1. Cytokinesis in the first two mitotic divisions

Cytokinesis in the first (top row) or second (bottom row) division in embryos expressing AIR-2::GFP (green, left, with H2B::mCherry and PH::mCherry in magenta) or NMY-2::GFP (green, right, with PH::mCherry in magenta). White arrowheads indicate first midbody that is internalized by AB and red arrowheads indicate the AB midbody, which is engulfed by EMS. Images are max Z projections of 15 central planes spaced 1μm apart taken every 90 seconds. Playback rate is 2 frames/second.

### Video S2. Lattice Light Sheet Imaging of Intestinal Cytokinesis

E8-E16 intestinal cell divisions imaged in embryos expressing AIR-2::GFP (green) with PH::mCherry (magenta) with lattice light sheet microscopy. Max intensity projection of 100 z-planes spaced 0.3 μm apart, acquired every 85 seconds.

### Video S3. Cytokinesis in the intestine epithelia

E8-E16 intestinal cell division in embryos expressing AIR-2::GFP (green, left, PH::mCherry in magenta), NMY-2::GFP (green, middle, tubulin::mCherry in magenta) or ZEN-4::GFP (green, right, tubulin::mCherry in magenta). The midbodies (indicated by arrowheads) form and move toward the apical surface. ZEN-4::GFP and NMY-2::GFP rapidly disappear, while AIR-2::GFP persists at the apical midline. Maximum z projections of 10 planes spaced 1μm apart, taken every 60 seconds. Playback rate is 6 frames/second.

### Video S4. High temporal resolution of midbody dynamics in the intestine

Imaging Earp cell division with high temporal resolution in an embryo expressing AIR-2::GFP (green, PH::mCherry in magenta) shows the lengthening of the central spindle and midbody migration event. Single plane images were acquired every 10 seconds. Playback rate is 15 frames/second.

### Video S5. Cytokinesis in E16 to E20 cell division

E16-E20 cell division in embryos expressing AIR-2::GFP (green) with PH::mCherry and H2B::mCherry (magenta). Max z projection of 15 z planes 1 μm apart that were acquired every 90 seconds. Playback rate is 6 frames/second.

### Video S6. Microtubule dynamics during E8-E16 in Aurora B mutants

WT (top row) and *air-2(or207)* (bottom row) mutant embryos expressing TBB-1::GFP (green in merge, right movie) and PH::mCherry and H2B::mCherry (middle, magenta in merge). Spindle midzone microtubules in WT embryos move to the midline where microtubules accumulate. In Aurora B mutant embryos, spindle midzone microtubules are diminished and cells that fail cytokinesis (right pair of gut cells) delay microtubule accumulation compared with the cells that do not fail. Max z projection of 10-15 z planes 1 μm apart that were acquired every 60 seconds. Playback rate is 6 frames/second.

### Video S7. Aurora B regulates adhesion dynamics during E8-E16 epithelial polarization

WT (top row) and *air-2(or207)* (bottom row) mutant embryos expressing HMP-1::GFP (green in merge, right movie) and TBB-1::mCherry (top middle, magenta in upper left merge) or PH::mCherry and H2B::mCherry (bottom middle, magenta in lower left merge). In WT, HMP-1::GFP accumulates at the furrow and midbody as it migrates to the apical surface where it accumulates during polarization. In Aurora B mutant embryos, HMP-1::GFP is reduced in the furrow and midbody and cells that fail cytokinesis (left pair of gut cells) delay adhesion accumulation at the midline. Max z projection of 10-15 z planes 1 μm apart that were acquired every 60 seconds. Playback rate is 6 frames/second.

### Video S8. Cytokinesis in the pharynx from ventral views

Cell division in pharyngeal precursor cells in embryos expressing AIR-2::GFP (green, left, H2B::mCherry in magenta), ZEN-4::GFP (green, middle left, TBB-1::mCherry in magenta), HMP-1::GFP (green, middle right, TBB-1::mCherry in magenta) or NMY-2::GFP (green, right, TBB-1::mCherry in magenta). Midbodies (white arrowheads) migrate toward pharyngeal midline after forming centrally between daughter cell pairs. Max z projection of 10-15 z planes 1 μm apart that were acquired every 90 seconds. Playback rate is 6 frames/second.

### Video S9. Lattice light sheet imaging of pharyngeal precursor cell divisions

Cytokinesis in pharyngeal precursor cell divisions in embryos expressing AIR-2::GFP (green, PH::mCherry in magenta). Max intensity projection of 101 z-planes spaced 0.35 μm apart, acquired every 60 seconds. Playback rate is 7 frames/second.

### Video S10. Cytokinesis in the sensilla dendrite development

Cell division in sensilla precursor cell divisions in embryos expressing AIR-2::GFP (green, left, H2B::mCherry in magenta), ZEN-4::GFP (green, middle left, TBB-1::mCherry in magenta), NMY-2::GFP (green, middle right, TBB-1::mCherry in magenta), or HMP-1::GFP (green, right, TBB-1::mCherry in magenta). Midbodies (white arrowheads) migrate into an apical cluster. Max z projection of 10 z planes 1 μm apart that were acquired every 90 seconds. Playback rate is 6 frames/second.

### Video S11. Lattice Light Sheet imaging of Sensilla Precursor Cell Division

Embryos expressing midbody flank marker AIR-2::GFP (green) with PH::mCherry (magenta) were imaged during the sensilla precursor divisions. Max intensity projection of 111 z-planes spaced 0.35 μm apart, acquired every 60 seconds. Playback rate is 7 frames/second.

### Video S12. Dendrite Extension of sensilla neurons

Embryo expressing AIR-2::GFP from the apical cluster stage of amphid polarization through the process of dendrite extension. Max z projection of 7 z planes 1 μm apart viewed from the ventral aspect and acquired every 90 seconds. Playback rate is 6 frames/second.

